# Local translation is engaged to sustain synaptic function in impaired Wallerian degeneration

**DOI:** 10.1101/2023.08.18.553870

**Authors:** Maria Paglione, Leonardo Restivo, Sarah Zakhia, Arnau Llobet Rosell, Marco Terenzio, Lukas J. Neukomm

## Abstract

After injury, the severed axon separated from the soma activates programmed axon degeneration, an evolutionarily conserved pathway to initiate its degeneration within a day. Conversely, severed projections deficient in programmed axon degeneration remain morphologically preserved with functional synapses for weeks to months after injury. How this synaptic function is sustained remains currently unknown. Here, we demonstrate that dNmnat-mediated over-expression attenuates programmed axon degeneration in distinct neuronal populations. Severed projections remain morphologically preserved for weeks after injury. When evoked, they elicit a postsynaptic behavior which is a readout for preserved synaptic function. We used ribosomal pulldown to isolate translatomes from these projections. Transcriptional profiling revealed several enriched biological classes. Identified candidates were validated in a screen using a novel system to automatically quantify evoked antennal grooming behavior as a proxy for preserved synaptic function. We used RNAi-mediated knockdown to identify mTOR as a mediator of local protein synthesis, and specifically candidates involved in protein ubiquitination and calcium homeostasis, required for preserved synaptic function. Our dataset uncovered several uncharacterized genes linked to human diseases. It may therefore offer insights into novel avenues for therapeutic treatments.

## Introduction

Wallerian degeneration is a simple and well-established system to investigate how damaged axons execute their destruction (Waller, 1850). Upon axonal injury (axotomy), the axon separated from the soma employs programmed axon degeneration to initiate its degeneration within a day. The resulting axonal debris is cleared by phagocytosis through neighboring glial cells in subsequent days (Raiders et al., 2021; Sapar & Han, 2019). Programmed axon degeneration is conserved among various species and appears to be hijacked in the absence of injury, in several neurological conditions (Coleman & Höke, 2020; Llobet Rosell & Neukomm, 2019).

In *Drosophila*, programmed axon degeneration is mediated by four genes and a single metabolite (Fang et al., 2012; Llobet Rosell et al., 2022; Neukomm et al., 2017; Osterloh et al., 2012; Paglione et al., 2020; Xiong et al., 2012). *Drosophila* nicotinamide mononucleotide adenylyltransferase (dNmnat) is synthesized in the soma and transported into the axon, where it is degraded by the E3 ubiquitin ligase Highwire (Hiw). Axonal transport and degradation result in steady-state dNmnat, which consumes nicotinamide mononucleotide (NMN) to generate nicotinamide adenine dinucleotide (NAD^+^) in an ATP-dependent manner. Upon injury, the axonal transport of dNmnat is abolished. Consequently, dNmnat rapidly decreases together with NMN consumption and NAD^+^ synthesis. The resulting increase of NMN activates *Drosophila* Sterile Alpha and TIR Motif-containing protein (dSarm), a NADase that pathologically depletes axonal NAD^+^. Low NAD^+^ results in the degeneration of the injured axon mediated by Axundead (Axed).

The manipulation of programmed axon degeneration results in severed axons and associated synapses (projections) that remain morphologically preserved for weeks to months. When evoked, they elicit a postsynaptic behavior, suggesting that synapses remain functionally preserved. In *hiw* mutant larvae, severed projections continue to elicit evoked excitatory junction potentials (EJPs) and spontaneous mini EJPs (mEJPs) in muscles up to 24 h after injury (Xiong et al., 2012). In diverse adult programmed axon degeneration mutants, the evoked stimulation of severed antennal grooming-inducing sensory neuron projections results in a behavior for at least two weeks after injury (Llobet Rosell et al., 2022; Neukomm et al., 2017; Paglione et al., 2020). This is also observed in mice, where muscle fibers respond to evoked severed motor projections for up to 5 days after injury (Mack et al., 2001). Thus, attenuated programmed axon degeneration preserves both morphology and synaptic function in severed projections for weeks after injury.

The metabolite NAD^+^ is instrumental in the morphological preservation of severed projections. While sustained NAD^+^ levels ensure preservation, its forced depletion triggers rapid axon and neurodegeneration (Essuman et al., 2017; Gerdts et al., 2015; Neukomm et al., 2017). However, the mechanisms that ensure the preservation of synaptic function are currently unknown.

In mice with attenuated programmed axon degeneration (Wallerian degeneration slow, *Wld^S^*), severed projections contain increased numbers of polyribosomes packed in multimembrane vesicles in the neurofilament space, suggesting that local protein synthesis may contribute to sustaining axonal homeostasis (Court et al., 2008). However, applying cycloheximide or emetine as protein synthesis inhibitors does not alter the morphological preservation after injury (Gilley & Coleman, 2010). Because local protein synthesis is crucial for synaptic plasticity (Holt et al., 2019; Ostroff et al., 2019; Yoon et al., 2012), we hypothesized that in severed projections, it contributes to the preservation of synaptic function.

Here, we use *Drosophila* to demonstrate that dNmnat-mediated over-expression (*dnmnat^OE^*) potently blocks programmed axon degeneration for weeks after injury. Consequently, severed projections of distinct sensory neuron populations remain morphologically preserved and their synapses functional. We employed tissue-specific ribosome pulldowns to isolate translated transcripts one week after injury. In-depth transcriptional profiling revealed several enriched biological classes. To validate our dataset, we established a novel system that automatically detects and quantifies antennal grooming behavior as a proxy for preserved synaptic function. We used this system to perform a high-throughput RNAi-mediated screen, which led to the identification of several protein ubiquitination and calcium homeostasis candidates as well as mTORC1-mediated local protein synthesis pathway. Our observations demonstrate that in a model of impaired Wallerian degeneration, local protein synthesis is required for the preservation of synaptic function.

## Results

### Preserved axonal morphology and synaptic function in severed projections with *dnmnat^OE^*-mediated attenuated programmed axon degeneration

We used a *Drosophila* model of impaired Wallerian degeneration to isolate and explore the axonal and synaptic translatome (Llobet Rosell & Neukomm, 2019). To attenuate programmed axon degeneration and isolate the local translatome, we decided against programmed axon degeneration mutants which are homozygous lethal. They require Mosaic Analyses with a Repressible Cell Marker (MARCM); thus the local translatome can only be isolated in sparse clonal neurons (Lee & Luo, 1999; Neukomm et al., 2014). We used the Gal4/UAS system to over-express dNmnat (*dnmnat^OE^*) in neuronal tissues. However, its over-expression in previous attempts only partially blocked programmed axon degeneration (MacDonald et al., 2006).

To test the *dnmant^OE^* preservation, we used an untagged cytoplasmatic *dnmnat* isoform in *dpr1^+^* sensory neuron clones in the wing (see Genotype List for details) (Zhai et al., 2006). In 7-day-old animals, one wing was subjected to partial injury, while the other served as an uninjured control. We quantified GFP-labeled injured and uninjured control axons at 20 and 25 °C, 7 and 14 days post axotomy (dpa). While wild-type axons degenerated within days after injury, robust *dnmnat^OE^*-mediated preservation was observed up to 14 dpa (Figure 1A, B). The preservation appeared slightly better at 25 °C; we thus conducted subsequent experiments at this temperature.

**Figure 1.**
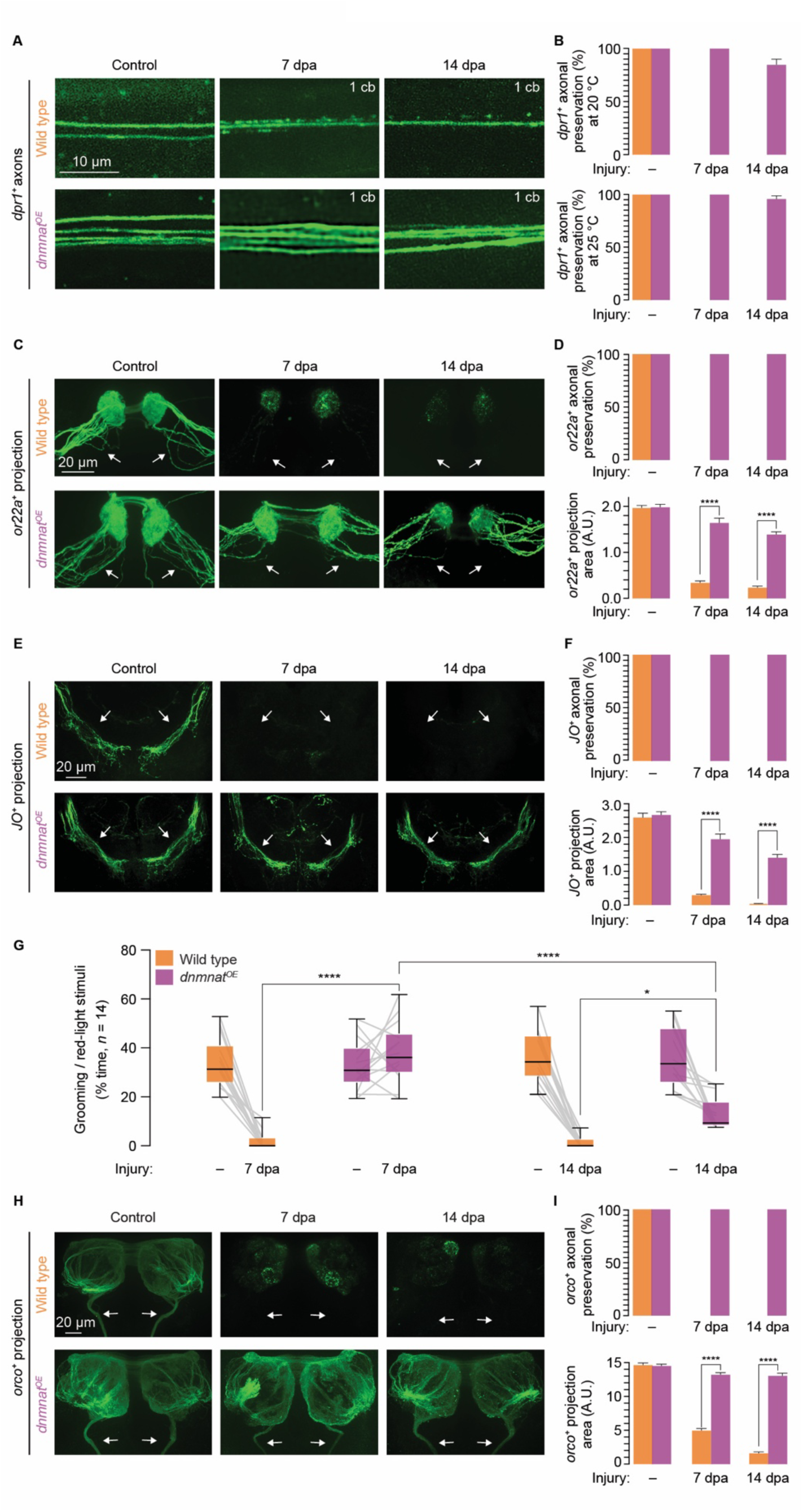
*dnmnat^OE^*-mediated attenuated programmed axon degeneration preserves axons and synapses for weeks after injury. **A** Examples of *drp1^+^* GFP-labeled wing axonal projections. Wild type and over-expression of dNmnat (*dnmnat^OE^*), top and bottom; uninjured (control), and after injury (7 and 14 days post axotomy (dpa)), respectively. Scale bar, 10 µm. The number of remaining cell bodies (cb), and therefore intact axons, is indicated in the upper right corner of each example. **B** Quantification of intact *drp1^+^* axons at 20 and 25 °C; top and bottom, respectively. Data, average ± SEM (*n* = 15 wings). **C** Examples of *or22a^+^* GFP-labeled olfactory receptor neuron (ORN) axonal projections (arrow). Wild type and *dnmnat^OE^*, top and bottom; control, and 7 and 14 dpa, respectively. Scale bar, 20 µm. **D** Quantification of intact *or22a^+^* axons and total axonal and synaptic area; top and bottom, respectively. Data, average ± SEM (*n* ≥ 10 brains). Unpaired two-tailed t-student test. **E** Examples of GFP-labeled *JO^+^* mechanosensory axonal projections. Wild type and *dnmnat^OE^*; control, and 7 and 14 dpa, respectively. Scale bar, 20 μm. **F** Quantification of intact *JO^+^* axons and total axonal and synaptic area. Data, average ± SEM (*n* ≥ 10 brains). Unpaired two-tailed t-student test. **G** Robust preservation of synaptic function in severed *dnmnat^OE^* projections at 7 dpa. Quantification of manually scored optogenetically-evoked antennal grooming behavior in uninjured and injured animals. Data, % time of red-light stimuli (*n* = 14 animals). Three-way ANOVA with Tukey’s multiple comparisons test. **H** Examples of olfactory receptor co-receptor (*orco^+^*) GFP-labeled wild-type and *dnmnat^OE^* projections. Wild type and *dnmnat^OE^*; control, and 7 and 14 dpa, respectively. Scale bar, 20 μm. **I** Quantification of *orco^+^* intact axons and projection area, respectively. Data, average ± SEM (*n* ≥ 5 brains). Unpaired two-tailed t-student test. ****, *p* < 0.0001; *, *p* < 0.05.

Next, we used *dnmnat^OE^* in approximately 20 *or22a^+^* GFP-labelled olfactory receptor neurons (ORN). Their cell bodies are housed in 3^rd^ antennal segments and project their axons into the antennal lobe (Vosshall et al., 2000). The bilateral ablation of 3^rd^ antennal segments results in the removal of the cell bodies and the degeneration of ipsi- and contralateral projections (MacDonald et al., 2006). While wild-type axons degenerated after injury, severed *dnmnat^OE^* projections remained strongly preserved at 7 and 14 dpa (Figure 1C). Quantifying *or22a^+^* axons and their projection area as a proxy for the synaptic field confirmed the strong *dnmnat^OE^*-mediated preservation (Figure 1D). We made similar observations with approximately 40 *JO^+^* mechano-sensory neurons in the Johnston’s Organ (JO). *JO^+^* cell bodies are housed in 2^nd^ antennal segments and project their axons into the subesophageal zone (Hampel et al., 2015, 2017). While the bilateral ablation of 2^nd^ antennal segments resulted in the complete degeneration of *JO^+^* wild-type projections, *dnmnat^OE^* projections remained preserved at 14 dpa (Figure 1E, F). Our observations suggest that *dnmnat^OE^* potently blocks the activation of programmed axon degeneration signaling, thereby preserving the morphology of severed projections in various neurons for weeks after injury.

We used the *JO^+^* neurons, which are required and sufficient for antennal grooming, to assess the preservation of synaptic function (Hampel et al., 2015, 2017; Neukomm et al., 2017; Paglione et al., 2020). First, we determined the age of adult animals where evoked antennal grooming behavior is consistently robust. Animals were fed with *all*-trans retinal during development, and adults expressing a red-shifted Channelrhodopsin (CsChrimson) in *JO^+^* neurons were subjected to three consecutive 10 s 10 Hz red-light exposures (see Material and Methods for details). Independent groups of adult animals were exposed once to red light from 0-1 to 7-8 days post eclosion (dpe). We observed a robust evoked antennal grooming behavior at 7-8 dpe and used this age for subsequent experiments (Supplementary Figure 1).

We applied the same injury paradigm in manipulated *JO^+^* neurons described above. While optogenetics failed to evoke antennal grooming in wild-type animals at 7 dpa, animals with *dnmnat^OE^ JO^+^* neurons continued to elicit antennal grooming (Figure 1G). We made a similar observation at 14 dpa, albeit with a significantly reduced evoked behavior (Figure 1G). Our observation strongly supports that in severed *dnmnat^OE^* projections, synapses remain functional for at least one week after injury.

The abundance of protein synthesis is lower in dendrites and axons than in the soma (Glock et al., 2021). To increase the biological material, we sought a Gal4 driver that labels an extended set of sensory neurons whose cell bodies can be readily ablated. We identified the pan-ORN Gal4 driver olfactory receptor co-receptor (*orco– Gal4*) that is transcriptionally active in approximately 800 ORNs (Vosshall et al., 2000; Wang et al., 2003). *orco^+^* cell bodies are housed in antennae and maxillary palps, and their axons project into the antennal lobe. Bilateral antennal and maxillary palp ablation resulted in the degeneration and debris clearance of wild-type axons at 14 dpa. In contrast, severed *dnmnat^OE^* projections remained extensively preserved (Figure 1H, I). These results demonstrate that *dnmnat^OE^* prevents axonal and synaptic degeneration for weeks after injury, regardless of the number of manipulated neurons. In subsequent experiments, we focused on 7 dpa to isolate the local translatome from severed projections, as morphology and synaptic function remained unchanged.

### Local axonal and synaptic translatomes of severed, preserved projections one week after injury

We sought to test whether the GFP-tagged large ribosomal subunit protein RpL10Ab (GFP::RpL10Ab) can be detected and thus used for translating ribosome affinity purification (TRAP) in severed projections (Thomas et al., 2012). While the Gal4/UAS-mediated expression of GFP::RpL10Ab was neither detected in 20 *or22a^+^* nor 40 *JO^+^* neurons, 800 *orco^+^* neurons revealed a GFP signal in Western blots derived from heads (Supplementary Figure 2A, B). Thus, *orco^+^* neurons were used to perform tissue-specific TRAP (Figure 2A). Briefly, GFP::RpL10Ab was expressed in *orco^+^* wild-type and *dnmnat^OE^* olfactory organs. Both genotypes were subjected to antennal and maxillary palp ablations, and at 7 dpa, 300 heads were collected and snap-frozen. Subsequently, axonal and synaptic-specific ribosomes/polysomes were immunoprecipitated with anti-GFP antibody-coated magnetic beads, mRNAs extracted, and reverse transcribed to establish cDNA libraries (Figure 2A, and Material and Methods).

**Figure 2.**
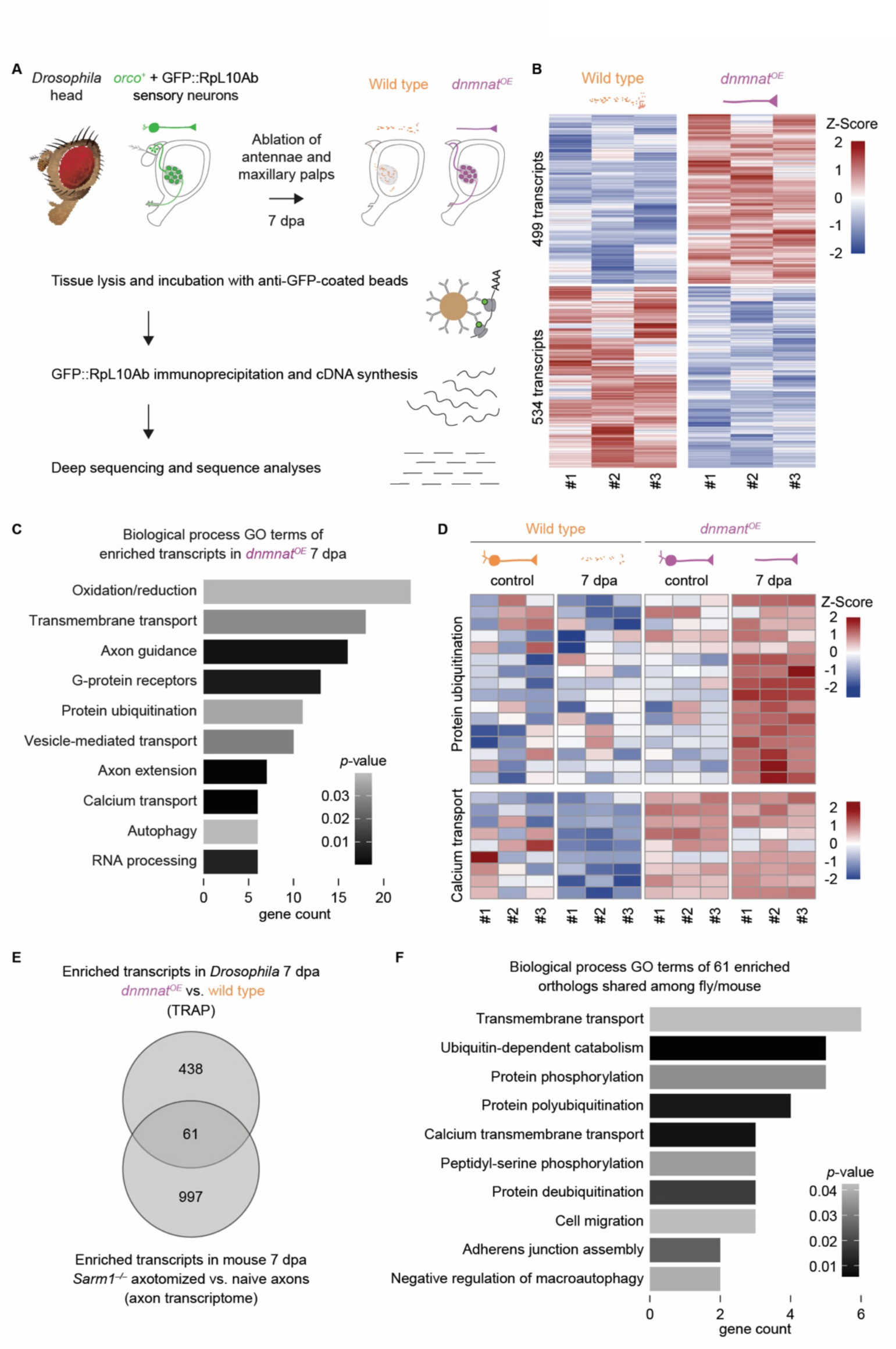
Identification of local translatomes in severed olfactory organ axons and synapses one week after injury. **A** Translating ribosome affinity purification (TRAP) as a strategy to isolate *orco^+^* translatomes from severed axons and their synapses 7 dpa. **B** Heat map of 1033 differentially enriched transcripts between wild type and *dnmnat^OE^* 7 dpa (Z-Score average). **C** Gene ontology (GO) term enrichment analysis (biological processes) of 499 *dnmnat^OE^*-enriched transcripts (fold change value > 0.6); sorted and plotted by *p*-value. **D** Heat map example of two significantly changed GO term groups, protein ubiquitination (*n* = 16) and calcium transport (*n* = 9), respectively. **E** Venn diagram of 499 *Drosophila dnmnat^OE^*-enriched TRAP transcripts and 1058 mouse *Sarm1^−/−^* KO transcripts (axonal transcriptome; axotomized versus naive axons (J. Jung et al., 2023)). **F** GO term analysis (biological processes) of 61 enriched fly/mouse ortholog transcripts.

We identified 1033 differentially enriched axonal and synaptic transcripts in the two genotypes at 7 dpa, with 499 transcripts enriched in *dnmnat^OE^* (Figure 2B). These enriched transcripts, with a regularized logarithm (rLog) fold change value > 0.6 and a false discovery rate (FDR) < 0.05, were run through the database for annotation, visualization, and integrated discovery (DAVID) for gene ontology (GO) analyses of biological processes (Figure 2C). Notably, we observed a significant transcript enrichment of calcium transport (*p* = 0.00017), axon extension (*p* = 0.0004), axon guidance (*p* = 0.002), RNA processing (*p* = 0.007), as well as oxidation/reduction, ubiquitination, and transmembrane transport (*p* = 0.03 each, respectively) (Figure 2C).

Primary cortical neurons lacking *Nmnat2* show an altered transcriptional profile compared to their controls (ZhenXian Niou, 2022). We therefore wondered whether *dnmnat^OE^* results in a similar, but likely inversed, change in the translatome profile in whole neurons. Consequently, we included uninjured controls of wild-type and *dnmnat^OE^* in our analyses. We identified transcripts of protein ubiquitination enriched in *dnmnat^OE^* solely after injury (Figure 2D). In contrast, transcripts of calcium transport were already enriched before injury, suggesting a potential response to *dnmnat^OE^* (Figure 2D). We made a similar observation with transcripts related to axon guidance and extension, G-protein receptors, transmembrane transport, vesicle-mediated transport, autophagy, RNA processing, and 50 % of the transcripts of oxidative stress (Supplementary Figure 3). Our analyses revealed enriched, *dnmnat^OE^*-independent and dependent transcripts of distinct biological GO terms.

A recent study examined *in vivo* mRNA decay in mammals by analyzing the axonal transcriptome in severed axons of *Sarm1^−/−^* mice (J. Jung et al., 2023). mRNAs were collected from *Sarm1^−/−^* retinal ganglion cell (RGC) axons at 7 dpa and compared to naïve (uninjured) axons of the same genotype. Notably, while *Sarm1* mutant mice have a different genotype than *dnmnat^OE^* flies, both genotypes prevent the activation of programmed axon degeneration after injury (Fang et al., 2012; Gilley & Coleman, 2010; Osterloh et al., 2012). The comparison of the two datasets (*Sarm1^−/−^* axonal transcriptome at 7 dpa vs. *dnmnat^OE^* TRAP translatome at 7 dpa) revealed 61 enriched, orthologous mouse/fly transcripts in both datasets (Figure 2E). Biological GO term analyses revealed an enrichment of transcripts involved in protein polyubiquitination (*p =* 0.009), calcium transmembrane transport (*p =* 0.008), and transmembrane transport (*p =* 0.042) (Figure 2F). Therefore, our analysis of mouse and fly orthologs revealed a common set of enriched biological GO term transcripts after axonal injury.

Next, we implemented an additional control using *orco–Gal4* alone (background) to further restrict our analyses (Supplementary Figure 4A). Both wild-type and *dnmnat^OE^* samples were filtered against this background (Supplementary Figure 4B, C, D). The additional stringent filtering led to the identification of 165 transcripts in *dnmnat^OE^* at 7 dpa (Supplementary Figure 4E). However, the biological GO term analyses showed that most transcripts were similar to those found in the initial dataset (Supplementary Figure 4F), where protein ubiquitination and calcium transport appeared significantly enriched in the local translatome datasets.

### A robust automated and high-accuracy antennal grooming detection system as a readout for preserved synaptic function

We aimed to validate whether the identified candidate transcripts are required to preserve synaptic function in severed, preserved projections one week after injury. Given the large number of candidates, we decided to develop a system that rapidly detects antennal grooming behavior, since manual analysis is labor intensive.

We developed two neural networks that perform frame-wise scoring of the movement of forelegs on the antennae, e.g., antennal grooming behavior. Briefly, Network 1 identifies the head region of the animal and enables the reduction of the analysis on a small area of the recording (Figure 3A). Network 2 uses this area and overlays three frames to identify pixel-wise standard deviations to generate a compressed image for antennal grooming behavior detection (Material and Methods) (Figure 3A). The system was trained and validated to identify grooming or no grooming events in these compressed images, resulting in a frame-wise probability of antennal grooming (Figure 3A). After training, the system produced probability scores similar to the manual scores of antennal grooming made by the trained observer (Figure 3B).

**Figure 3.**
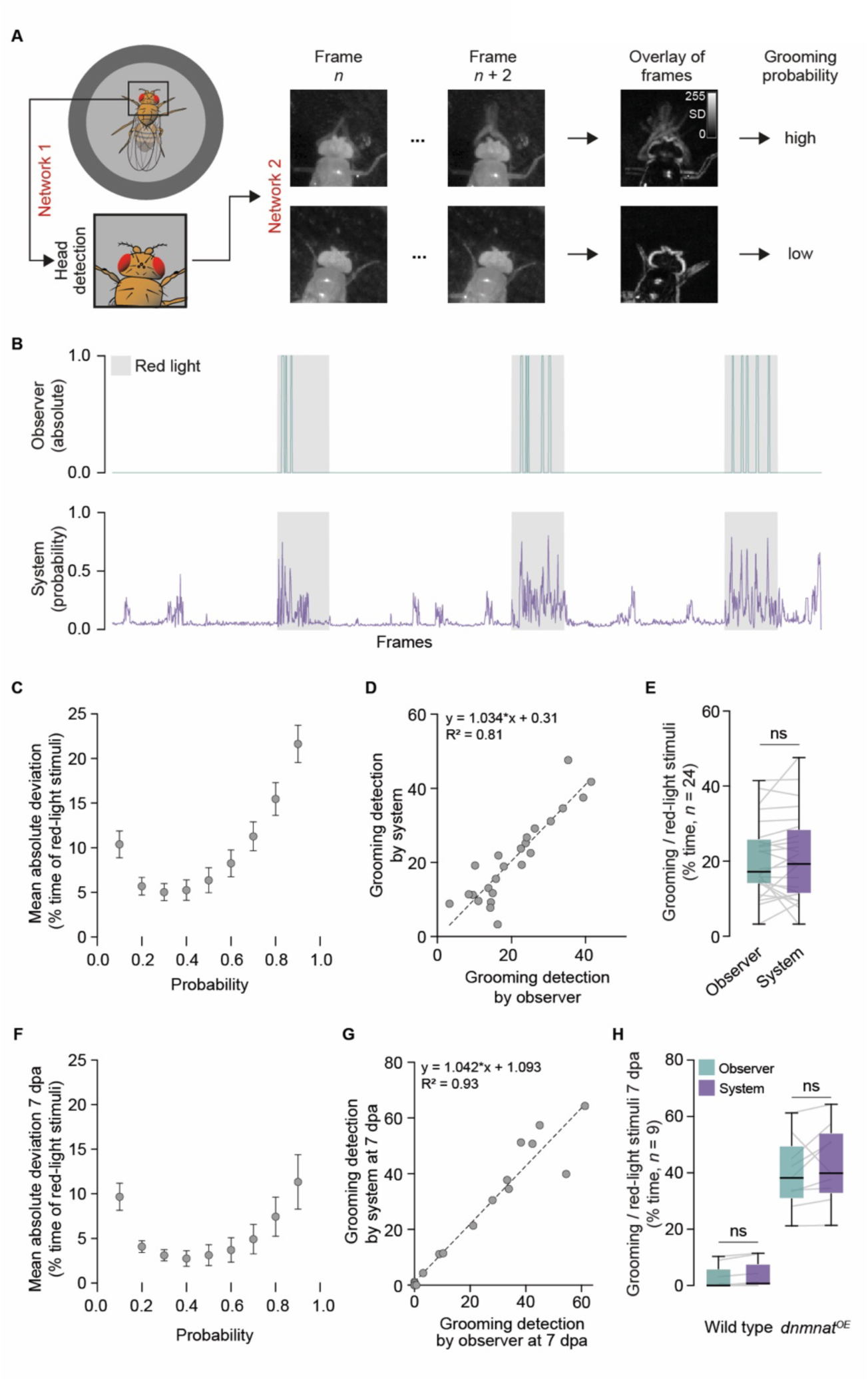
Automated high-accuracy antennal grooming behavior detection system. **A** Illustration of the two-network system for head and antennal grooming detection (networks 1 and 2, respectively). **B** Examples of antennal grooming scores by the observer (absolute) and the system (probability); green and purple, top and bottom, respectively. Grey, 10 s 10 Hz red-light stimulus. **C** Mean absolute deviation of grooming detection between observer and system (mean ± SEM, % time of red-light stimuli). **D** Linear regression of grooming detection by system and observer (% time of red-light stimuli; probability ≥ 0.3; *n* = 24 animals). **E** Grooming detection quantified by observer and system (% time of red-light stimuli, *n* = 24 animals). Paired two-tailed t-student test. **F** Mean absolute deviation between manual and automatic grooming detection after injury (7 dpa; mean ± SEM, % time of red-light stimuli). **G** Linear regression of grooming detection between observer and system at 7 dpa (% time of red-light stimuli; probability ≥ 0.4; *n* = 18 animals). **H** Grooming detection quantified by observer and system at 7 dpa (% time of red-light stimuli, *n* = 9 animals). Paired two-tailed t-student test; ns, *p* > 0.05.

To identify the probability cutoff where the quantification difference between the probability score assigned by the system and the manual score by the observer is minimal, the mean absolute differences were plotted in a probability-dependent manner (Supplementary Figure 5). Once the probability threshold was determined, the system was further validated by a test set. A minimal difference was observed with a probability ≥ 0.3 (Figure 3C). A linear regression between system and observer scores confirmed a strong correlation with a ≥ 0.3 probability cutoff (Figure 3D). Despite the > 93 % performance achieved by the system on the test set (Material and Methods, Supplementary Figure 5), we further compared the detection of antennal grooming from the system with the scores from the observer on the test set. Grooming detection was not statistically significant between the system and the observer (Figure 3E). We therefore conclude that our newly established two-network system plots accurate probabilities of evoked antennal grooming (Video 1).

We employed this system primarily after antennal ablation. However, we observed that even the presence or absence of antennae has a significant impact on the performance of network 2. We used a second, different training set comprised of videos where antennae were removed (e.g., at 7 dpa) to identify grooming or no-grooming events in compressed images after injury. We repeated the same procedure with ablated antennae and determined a ≥ 0.4 probability cutoff where the difference between the system and the manual scores was minimal (Figure 3F). A linear regression between system and observer scores revealed a similar strong correlation (Figure 3G). We used newly recorded videos where antennal grooming detection was compared between wild type and *dnmnat^OE^* at 7 dpa with no statistical difference between the system and the observer (Figure 3H), confirming the accuracy of the predictions (Video 2, 3).

To validate the accuracy of the system, we compared the grooming detection output of the system with manual scores obtained by two independent observers in uninjured controls and at 7 dpa. The result of the system was in line with both observers. It suggests the system produces scores in the range of the variability measured among observers (Supplementary Figure 6A, B).

We also tested whether different experimental conditions affect grooming behavior picked up by the observer or system. We combined two distinct temperatures (20 and 25 °C) with red-light intensities (8 and 14 mW/cm^2^). While red-light intensities did not alter the behavior, temperature-dependent nuances were identified by both the system and the observer. Animals groomed significantly more at 25 °C (Supplementary Figure 7). Thus, the system reliably detects the behavior of evoked antennal grooming and can be used to perform unbiased screens to identify subtle behavioral changes.

### RNAi-mediated knockdown of protein ubiquitination and Ca^2+^ transport candidates alter preserved synaptic function

We used automated grooming detection to validate candidates from our translatome dataset for preserved synaptic function after injury. Highly translated mRNAs are selectively degraded in axons; thus their pools decrease over time (J. Jung et al., 2023). In our paradigm, while uninjured and injured *dnmnat^OE^* animals harbor a robust antennal grooming behavior, the specific candidate RNAi-mediated knockdown should reduce the behavior specifically at 7 dpa (Figure 4A).

**Figure 4.**
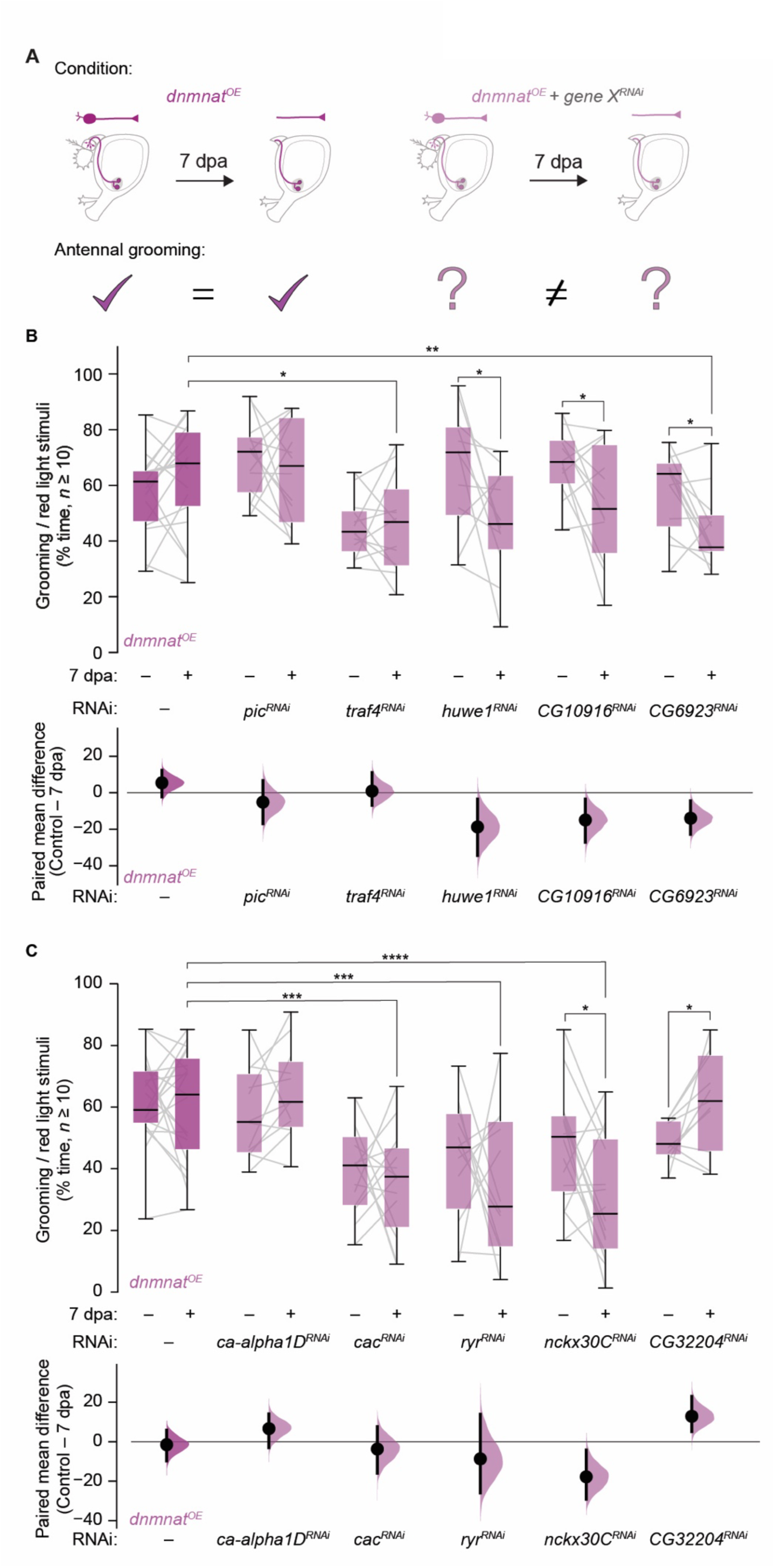
RNAi-mediated knockdown of candidates involved in protein ubiquitination and calcium transport required for preserved synaptic function one week after injury. **A** RNAi-based strategy to identify candidates required for synaptic function. **B** Top, screen of protein ubiquitination candidate genes. Grooming as % time of red-light stimuli. Bottom, paired mean difference as bootstrap sampling distribution. Mean differences, dots; vertical error bars, 95% confidence intervals, respectively. **C** Top, screen of calcium transport candidate genes. Bottom, paired mean difference. Data, % time of red-light stimuli (*n* ≥ 10 animals). Paired two-tailed t-student and Two-Way ANOVA with Dunnett’s multiple comparisons test; ****, *p* < 0.0001; ***, *p* < 0.001; **, *p* < 0.01; *, *p* < 0.05.

The comparisons of the mammalian dataset with our background filtering strategy prompted us to validate protein ubiquitination and calcium transport GO term classes in our dataset (Figure 2E, F, Supplementary Figure 4E, F). In *dnmnat^OE^ JO^+^* neurons, we targeted 9 enriched protein ubiquitination candidates by RNAi in uninjured controls and at 7 dpa. We observed three major phenotypes. First, several candidate RNAi experiments did not change evoked antennal grooming behavior in uninjured controls or at 7 dpa, e.g., *pic^RNAi^*(Figure 4B and Supplementary Figure 8). Second, reduced grooming was observed in uninjured controls and at 7 dpa, e.g., *traf4^RNAi^* (Figure 4B). Third, grooming behavior was unaltered in uninjured controls but reduced at 7 dpa, e.g., *huwe1^RNAi^* and the uncharacterized coding gene 10916 (*CG10916^RNAi^*). RNAi-mediated knockdown of *CG6923* exhibited the most substantial decrease in evoked grooming behavior at 7 dpa (Figure 4B). Thus, our RNAi-mediated approach provides evidence that the local translation of various transcripts involved in protein ubiquitination is required to sustain *dnmnat^OE^*-mediated synaptic function one week after injury.

Perturbed axonal calcium homeostasis often results in axon degeneration (Stirling & Stys, 2010). Thus, we also tested 7 identified Ca^2+^ transport candidate transcripts by RNAi. We observed similar phenotypes as described above. In *JO^+^ dnmnat^OE^* neurons, knockdown of several candidates did not alter evoked antennal grooming behavior in uninjured controls and at 7 dpa, including the pore-forming alpha^1^ subunit D of the L-type voltage-gated Ca^2+^ channel (*ca-alpha1D^RNAi^*) (Figure 4C, and Supplementary Figure 8A). RNAi-mediated knockdown of *cacophony* (*cac^RNAi^*) reduced the behavior in uninjured controls and at 7 dpa (Figure 4C). Knockdown of the endoplasmatic reticulum ryanodine receptor (*ryr^RNAi^*) required for Ca^2+^ release from intra-axonal stores also decreased grooming in uninjured controls and at 7 dpa. In contrast, RNAi-mediated knockdown of the sodium/potassium/calcium exchanger *nckx30C* (*nckx30C^RNAi^*) resulted in a reduced behavior solely at 7 dpa (Figure 4C), as observed with two different RNAi constructs (Supplementary Figure 8B). We also identified *CG32204*, where *CG32204^RNAi^* increases grooming behavior at 7 dpa (Figure 4C). Our observation provides evidence that local translation of transcripts involved in calcium transport is crucial for *dnmnat^OE^*-mediated preservation of synaptic function.

The RNAi-based validation of the candidates identified by TRAP prompted us to ask whether a direct impairment of local translation alters synaptic function. We sought to test mTOR signaling, which is required for local axonal translation following nerve injury (Terenzio et al., 2018). Raptor is a component of the mTOR complex 1 (mTORC1) and regulates protein synthesis through ribosomal protein S6 kinase (S6k) (Figure 5A). Notably, *raptor* was enriched at 7 dpa in our dataset (Figure 2B). We therefore targeted *raptor* and *s6k* by RNAi. In *JO^+^ dnmnat^OE^* neurons, RNAi-mediated knockdown of *raptor* (*raptor^RNAi^*) resulted in a decreased behavior in uninjured controls injury, and at 7 dpa, we observed a statistically non-significant trend of a further decrease (Figure 5B). In contrast, *s6k^RNAi^* harbored an impaired evoked antennal grooming behavior at 7 dpa (Figure 5B). Our observations support our TRAP-mediated identification of transcripts locally translated in severed axons and their synapses. These projections appear to engage mTOR signaling-mediated local protein synthesis to ensure ubiquitin-mediated protein degradation, to sustain amino acids for translation, and combat calcium imbalance.

**Figure 5.**
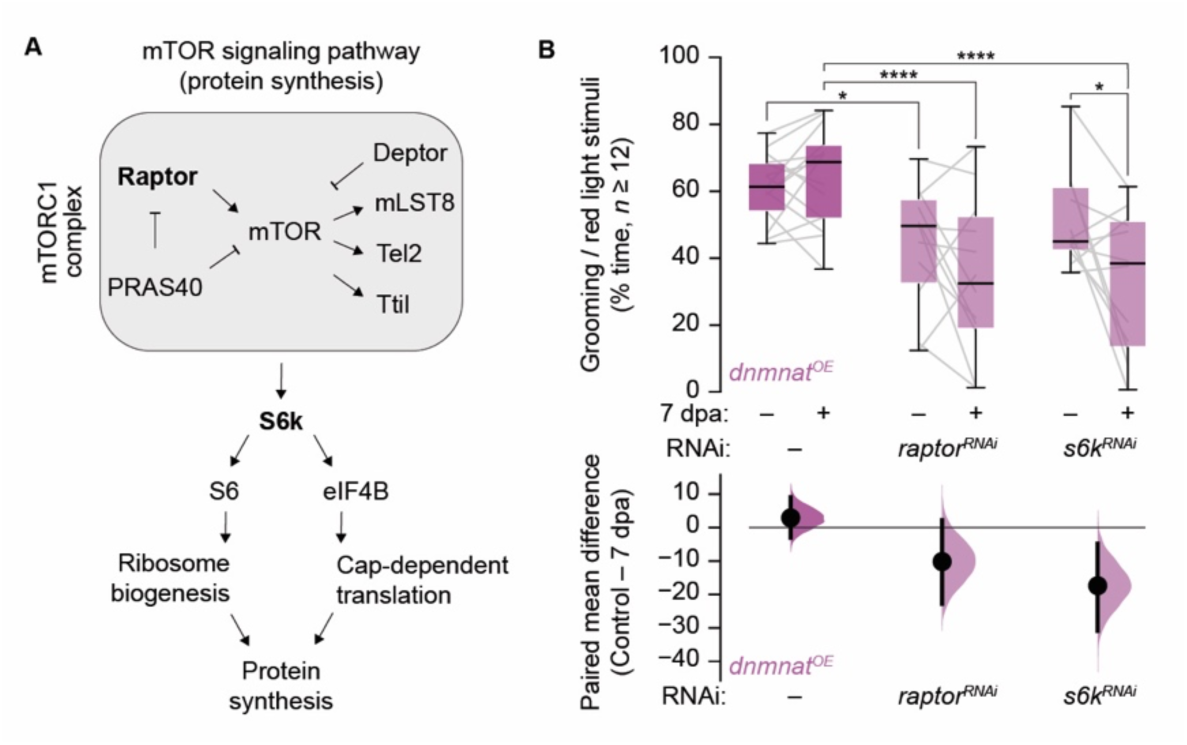
RNAi-mediated knockdown of mTOR signaling candidates reduces preserved synaptic function one week after injury. **A** Protein synthesis mediated by the mTOR signaling pathway. Bold, candidates tested by RNAi. **B** RNAi targeting *raptor* and *s6k* result in reduced antennal grooming after injury. Top, Grooming as % time of red-light stimuli; bottom, paired mean difference as bootstrap sampling distribution. Data, % time of red-light stimuli (*n* ≥ 12 animals). Paired two-tailed t-student test and Two-Way ANOVA with Dunnett’s multiple comparisons test; ****, *p* < 0.0001; *, *p* < 0.05.

## Discussion

Here, we investigated how severed projections, with attenuated programmed axon degeneration, employ local protein synthesis to sustain synaptic function for at least one week after injury. We over-expressed *Drosophila* Nmnat (*dnmnat^OE^*) to attenuate programmed axon degeneration, thus preserving both axonal morphology and synaptic function after injury. dNmnat/NMNAT2 is evolutionarily conserved and consumes NMN as a substrate to generate NAD^+^ in an ATP-dependent manner (Brazill et al., 2017; Llobet Rosell et al., 2022; Zhai et al., 2006, 2009).

The discovery of the Wallerian degeneration slow (*Wld^S^*) mouse provides the basis for our current understanding of the preserving NMNAT function (Lunn et al., 1989). In *Wld^S^*, a complex genomic rearrangement led to the fusion of the N-terminal 70 amino acids of UBE4b and full-length NMNAT1, referred to as WLD^S^ (Mack et al., 2001). The *Wld^S^* coding gene is inserted as a triplication between *Ube4b* and *Nmnat1.* It results in over-expressed WLD^S^, which is relocated from the nucleus to the axon. After injury, WLD^S^ persists in the severed axon, while endogenous NMNAT2 is rapidly degraded (Gilley & Coleman, 2010), resulting in preserved axonal morphology and synaptic function (Mack et al., 2001). Various studies confirmed the morphological preservation mediated by dNmnat in *Drosophila* (Fang et al., 2012; Llobet Rosell et al., 2022; MacDonald et al., 2006). Therefore, high levels or degradation-insensitive variants of dNmnat/NMNAT are a powerful tool for attenuating programmed axon degeneration and studying how severed projections remain preserved.

Here, we used an untagged dNmnat (Zhai et al., 2006) and observed that the morphological preservation lasts for at least two weeks in multiple neurons. It contrasts previous observations where the over-expression of N-terminal Myc-tagged dNmnat preserved axonal morphology for approximately 5 days after injury. (MacDonald et al., 2006). This enhanced neuroprotection may be attributed to untagged dNmnat or different expression levels due to distinct transgene insertion sites.

After injury, in the soma-attached proximal axon, axonal regeneration is initiated through mTOR-mediated local axonal translation (Terenzio et al., 2018). More broadly, mTORC1 signaling mediates translation and tissue regeneration in axolotl compared to non-regenerative tissue in mice (Zhulyn et al., 2023). Surprisingly, severed distal *Wld^S^* projections also harbor increased numbers of polyribosomes, suggesting that local protein synthesis may be in place to exert preservation (Court et al., 2008). We therefore hypothesized that severed projections engage maintenance through local protein synthesis. This appears in stark contrast with preserved *Wld^S^* axonal morphology that does not depend on local translation (Gilley & Coleman, 2010). Nevertheless, we revisited this question, but in the context of preserved synaptic function.

Local mRNA translation in axons and synapses is less abundant than in cell bodies (Glock et al., 2021). To increase the biological tissue for TRAP, we identified *orco– Gal4*, which labels 800 olfactory receptor neurons with cell bodies housed in antennae and maxillary palps (Larsson et al., 2004). Their cell bodies can be readily ablated without killing the animals. However, we observed an increase in mortality in wild type but not in animals with preserved projections at 14 dpa (e.g., *orco^+^ dnmnat^OE^*). To avoid the loss of animals, we used 7 dpa to perform TRAP and translatome analyses. Our study identified around 500 enriched transcripts by TRAP in severed, preserved projections of adult flies at 7 days after injury. Many transcripts are associated with oxidative/reduction processes, a response known to be activated after injury to counteract the increased production of reactive oxygen species (ROS) (Llobet Rosell & Neukomm, 2019). Furthermore, we identified vesicle-mediated transport transcripts to facilitate pre- and postsynaptic communication. Additionally, we isolated transcripts associated with RNA processing. RNA binding proteins regulate mRNA axonal transport and local translation (Ederle & Dormann, 2017; Ishiguro et al., 2016; Thelen & Kye, 2020). This could suggest that translationally silent mRNAs may be stored locally and utilized to produce multiple copies of a protein when needed, providing an efficient response mechanism in emergencies (H. Jung et al., 2012). Among them was the RNA-binding protein 4F (Rnp4F), encoding an evolutionarily conserved RNA-binding protein. We also identified uncharacterized genes, *CG32533* and *CG1582*, which are predicted to possess RNA binding and helicase activity. Our findings thus add complexity to RNA processing and local translation.

In addition, our analysis revealed two types of transcriptional enrichments. We examined the transcripts in preserved projections compared to their wild-type controls, and also considered the genetic background (e.g., wild type vs. *dnmnat^OE^*). One GO-term class of transcripts, protein polyubiquitination, was enriched exclusively in severed preserved *dnmnat^OE^* projections, as a marker for protein degradation. This increase suggests that protein degradation, generally in balance with protein synthesis, provides new amino acids for local translation (Ding et al., 2007; Jarome & Helmstetter, 2014). This enrichment indicates a response to the actual injury, which is independent of the genetic background. In line with our observation, a recent study used *Sarm1^−/−^* knockout axons to investigate mRNA decay, resulting in a comprehensive transcriptomic profile (Jung et al., 2023). The comparison of both datasets revealed a common significant enrichment of transcripts associated with protein ubiquitination. Thus local translation could be activated as a response to injury in preserved projections. In contrast, all other GO-term classes were enriched in uninjured *dnmnat^OE^* controls, suggesting that *dnmnat^OE^* changes the neuronal transcriptome.

Combining identified conserved mouse/fly transcripts with a more stringent filtering approach revealed an enrichment of transcripts related to calcium transport/homeostasis. A conditional knockout (cKO) of *Nmnat2* in cortical glutamatergic neurons supports our observations, where transcriptomic analyses revealed that NMNAT2 loss results in a significantly reduced Ca^2+^ transport/homeostasis (ZhenXian Niou, 2022). In line with our observations, elevated calcium levels can activate calpain proteases to execute axon degeneration (Yang et al., 2015). Intriguingly, voltage-dependent sodium channels are among the targets of calpains. The channel degradation leads to increased axonal calcium and subsequent axon degeneration (Iwata et al., 2004; Llobet Rosell & Neukomm, 2019; Rishal & Fainzilber, 2014).

Identified candidates and their subsequent validation led us to speculate that protein synthesis forms the basis for preserved synaptic function. Since we observed increased *raptor* transcripts in *dnmnat^OE^* at 7 dpa, we conducted knockdown experiments targeting Raptor and S6K, mediators of the mTORC1 pathway. The specific reduction of evoked antennal grooming behavior after injury with *s6k^RNAi^* suggests an intricate involvement of the mTORC1 pathway in regulating preserved synaptic function through local translation.

Based on our observations, we propose that severed projections with attenuated programmed axon degeneration employ local protein synthesis through axonal/synaptic polyribosomes. Among the translated transcripts are candidates that help to cope with the turnover of already translated polypeptides by polyubiquitination and calcium buffering. This model is supported by the impairment of either local translation or calcium transport and polyubiquitination candidates, resulting in reduced synaptic function. Altogether, our findings support the hypothesis of the “autonomous axon” (Alvarez, 2001), where local protein synthesis, with a pool of mRNAs, ensures the continued functional adjustments of projections where a nucleus is far away, or soma cut off.

Interestingly, axons degenerate within a few days in species such as flies and mice, while they persist for weeks to months in various invertebrates and some vertebrates (Bittner, 1988, 1991). Under specific circumstances, programmed axon degeneration is beneficial. For example, it may serve as a first line of defense against viral infection (Sundaramoorthy et al., 2020). Rabies viruses infect peripheral nerve endings, hijack axonal transport, and travel from the periphery to the central nervous system (Schnell et al., 2010). Here, engaged programmed axon degeneration impedes the spread of rabies among interconnected neurons *ex vivo* (Sundaramoorthy et al., 2020).

In contrast, programmed axon degeneration may be disadvantageous during long cold exposures. In humans, exposure to prolonged cold temperature results in damaged peripheral nerves, as observed in soldiers during World War II (Denny-Brown et al., 1945; Schaumburg et al., 1967). During winter, in warm-blooded hibernating vertebrates such as ground squirrels, the body temperature falls from 38 to 5 °C (Albuquerque et al., 1978). Cold exposures may result in partially damaged peripheral nerves, which is also observed in chemotherapy-induced peripheral neuropathy (Coleman & Höke, 2020). Here, engaged programmed axon degeneration is detrimental for the host, whereas its absence or attenuation appears beneficial.

The above examples demonstrate that programmed axon degeneration is a double-edged sword. It must be tightly controlled and solely engaged under specific circumstances. In some species, may be completely absent due to selective evolutionary pressure. Here, local translation ensures sustained maintenance of synaptic plasticity, which we observe in our model of impaired Wallerian degeneration.

Our comprehensive dataset revealed different uncharacterized genes in *Drosophila* linked to human diseases associated with impaired axonal transport and local translation. *CG13531* is an enigmatic gene with implications in axon extension and synaptic vesicle transport. Human ortholog(s) are predicted to be linked to Charcot-Marie-Tooth type 2X, amyotrophic lateral sclerosis type 5, and hereditary spastic paraplegia 11 (Yamaguchi et al., 2021; Yamaguchi & Takashima, 2018; Zhang et al., 2018). These observations highlight Wallerian degeneration, and the fruit fly, as powerful systems to gain further insights into human disease. Identifying and characterizing conserved fly/mammalian genes may therefore provide novel therapeutic avenues to slow or halt human axonopathies.

## Supporting information

Genotype list

## Acknowledgments

We would like to thank Dr. Takakazu Yokokura for the access to and support in the OIST fly facility. This work was supported by a Swiss National Science Foundation (SNSF) Assistant Professor Starting Grant Awards (PP00P3_176855 and PP00P3_211015), the International Foundation for Research in Paraplegia (P180), and SNSF Spark (190919) to LJN; a Japanese Society for the Promotion of Science (JSPS) Fellowship (GR20107) to MP and research grant (23H02414) to MT.

## Material and Methods

### Husbandry and genetics

*Drosophila melanogaster* was used to perform experiments. Animals were raised on Nutri-Fly Bloomington Formulation food at 25 °C on a 12h:12h light-dark cycle unless otherwise specified. Genotypes are listed in the genotype list.

### Wing injury assay and axonal preservation analysis

#### Wing assay

Animals were aged for 5 - 10 days at 20 or 25 °C. Wing injuries were applied as previously described (Llobet Rosell et al., 2022; Paglione et al., 2020). Briefly, one wing of each anesthetized animal was cut with micro scissors roughly in the middle, and was returned to an individual vial.

#### Axonal preservation analysis

The distal, cut-off wing was mounted on a microscopy slide in Halocarbon Oil 27 and covered with a coverslip. The number of neuronal cell body clones in the cut-off wing was counted with an epifluorescence microscope. It represents the number of injured axons. The second wing was kept as uninjured control for subsequent analysis. To count intact or degenerated axons at 7 and 14 days post-axotomy (dpa), the injured and uninjured control wings were mounted and imaged using a spinning disk microscope.

### Antennal and maxillary palp ablation, brain dissection, immunohistochemistry, image acquisition and analysis

#### Antennal or antennal and maxillary palp ablation

Animals were aged for 5 - 10 days at 25 °C. Anesthetized animals were subjected to bilateral antennal ablation or bilateral antennal and maxillary palp ablation with high precision and ultra-fine tweezers (Paglione et al., 2020). Animals were recovered in food-containing vials for a specified time.

#### Brain dissection

Adult brains were dissected at 7 and 14 dpa with the corresponding uninjured control, using a modification of a previously described protocol (Paglione et al., 2020).

#### Immunohistochemistry

Decapitated adult heads were pre-fixed in 4 % paraformaldehyde (PFA) in 0.1 % Triton X-100 in 1x phosphate-buffered saline (PBS), pH 7.4 (PTX) for 20 min at room temperature (RT), and subsequently washed in PTX for 5 × 2 min. The dissection of the brains was performed in PTX, followed by brain fixation in 4 % formaldehyde in PTX for 10 min at RT. After fixation, the brains were washed in PTX for 5 × 2 min and blocked in 10 % normal goat serum (Jackson Immuno) in PTX (blocking solution) for 1 h at RT. Primary antibody incubation was performed in a blocking solution containing 1:1000 chicken anti-GFP (Rockland) overnight at 4 °C. Brains were washed with PTX for 3 × 10 min at RT and incubated with secondary antibodies in PTX (1:200 goat anti-chicken IgY H&L DyLight 488 (Abcam)) for 2 h at RT. Brains were washed with PTX for 3 × 10 min at RT and mounted in Vectashield.

#### Image acquisition and analysis

Images were acquired along the z-axis with a step size of 0.6 μm using a Nikon Spinning Disk confocal microscope. Image quantification was analyzed with ImageJ (NIH).

### Western blots

For each genotype, 20 adult heads were ground in 100 µl of Laemmli sample buffer (2 heads/10 µl) and heated for 10 min at 95 °C. 20 µl of the resulting lysate was loaded per well into 4-12 % SurePAGE^TM^ gels (GeneScript) with MOPS running buffer for higher molecular weight proteins. Protein separation was carried out at 80 V. Precision Plus Protein^TM^ Kaleidoscope^TM^ Prestained Protein Ladder (Bio-Rad) was used as a molecular weight marker. The proteins were wet-transferred onto PVDF membranes (Bio-Rad) at 110 V for 70 min at 4 °C. Membranes were washed with Tris-buffered saline (TBS) containing 0.1 % Tween^®^ 20 (Merk) (TBS-T) for 5 min at RT. Membranes were blocked with 5 % milk (Carl-Roth) in TBS-T for 1h at RT, and incubated in a blocking solution containing primary antibodies (1:5000 rabbit anti-GFP (Abcam), and 1:15000 mouse anti-Tubulin (Sigma)) over night at 4°C. Membranes were rinsed with TBS-T for 3 × 10 min and incubated with secondary antibodies in 5 % milk in TBS-T (1:10000 goat anti-rabbit IgG (H+L) Dylight 800, and 1:10000 goat anti-mouse IgG (H+L) Dylight 680 (ThermoFisher)) for 1 h at RT. Membranes were then rinsed with TBS-T for 3 × 10 min before signal acquisition. Fluorescent signals were acquired using Odissey® DLx (LI-COR), and image quantification was performed with ImageJ (NIH) through densitometric analysis.

### Translation Ribosome Affinity Purification (TRAP)

#### Preparation of samples

The ribosomal pulldown was performed by adapting the already described protocol (Rozenbaum et al., 2018). In brief, for each genotype and condition, 300 adult animals were collected, snap-frozen in liquid nitrogen and stored at −80 °C before processing. The frozen samples were homogenized using 2-3 % weight per volume of homogenization buffer (100 mM KCl, 12 mM MgCl_2_, 1 % NP-40, 50 mM Tris, pH 7.4) supplemented with β-mercaptoethanol (10 µl / ml buffer solution). The lysate was centrifuged at 10000 rpm for 10 min at 4 °C, and the supernatant was collected in a protein low-binding tube. 10 % was kept for the input and stored at −80 °C.

#### Preparation of beads

For each sample, 50 µl of magnetic agarose GFP-Trap beads were added to a RNase/DNase-free microcentrifuge tube. The beads were rinsed with homogenization buffer supplemented with 1 mM DTT, 150 µg/ml cycloheximide, 450 U/ml RNasin OUT, 10 mM ribonucleoside vanadyl complexes (RVC) and protease inhibitor (Sigma, P8340)).

#### GFP pulldown

The homogenized heads were incubated with magnetic agarose GFP-Trap beads to pulldown ribosomes with associated mRNAs at 4 °C over night. The beads were washed in high salt buffer (300 mM KCl, 12 mM MgCl_2_, 1 % NP-40, 1 mM DTT, 150 µg/ml Cycloheximide, 450 U/mL RNasin OUT, protease inhibitor (Sigma, P8340), 50 mM Tris, pH 7.4) 3 × 5 min at 4 °C. Immediately after removing the final high salt wash buffer, samples were placed in a magnet, and 350 µl of lysis buffer was added to the beads (LB, RNA plus Kit for the inputs or LB1, RNA Plus XS kit, for the pulldowns). RNA was extracted from ribosomes bound to the beads according to the NucleoSpin RNA (input) and RNA Plus XS (pulldowns) kit manufacturer’s recommendations.

### Library preparation, sequencing, and data analysis

The library preparation was performed with poly-A purification for input samples and skipped poly-A for the pulldowns, both without rRNA depletion. The libraries were sequenced on Pair End, using an Illumina NovaSeq 6000 SP Whole sequencer, with 150 bp determined. All reads were analyzed by a quality control check using FASTQC, followed by trimming with Trimmomatic (Bolger et al., 2014), where unpaired reads and adaptor sequences were removed based on the quality scores. Then, another control quality check was performed.

Trimmed high-quality reads were aligned to the *Drosophila* genome (*Drosophila melanogaster*/dmel_r6.43_FB2021_06) using Hisat2 (Kim et al., 2015). The alignment files were sorted, indexed, and converted with Samtools to count reads with Htseq-Count (Anders et al., 2014). Library normalization and statistical analysis of differentially expressed genes (DEG) were analyzed by DESeq2 (Love et al., 2014), considering an adjusted *p*-value ≤ 0.05 and regularized logarithm (rLog) fold change smaller than −0.6 (down-regulated) or greater than 0.6 (up-regulated).

Gene Ontology (GO) term enrichment analysis for biological processes was performed with the Database for Annotation, Visualization, and Integrated Discovery (DAVID) 6.8 (Huang et al., 2009). Statistical analyses and data visualization were conducted in R using the base, dplyr, pheatmap, and ggplot2 packages.

### Comparative analysis with published data set

For the comparative analysis with the mammalian data, transcripts with adjusted *p*-value ≤ 0.05 and log2 Fold Change > 0.6 (up-regulated) were selected from Table S6B (J. Jung et al., 2023). For rapid identification of orthologs, the list of the selected Ensembl Gene ID was analyzed with the *Drosophila* Integrative Ortholog Prediction Tool (DIOPT; http://www.flyrnai.org/diopt) (Hu et al., 2011). Only orthologs with high rank and best score were chosen to compare with our data set.

### Optogenetics-induced antennal grooming behavior

Genetic crosses were performed in the darkness on standard cornmeal food in aluminum-covered vials supplemented with 200 µM all-trans retinoic acid (Hampel et al., 2015; Paglione et al., 2020). Optogenetic experiments to induce antennal grooming were conducted in the dark. Animals were visualized and recorded using an 850 nm infrared light source at an intensity of 2 mW/cm2 (Mightex, Toronto, CA). For CsChrimson activation, a 656 nm red-light source (Mightex) was used at 8 or 14 mW/cm^2^ intensities. Animals were subjected to a 10 s 10 Hz red-light exposure, followed by a 30 s recovery phase where the red light was turned off. This repetition cycle was performed three times in total. Red light stimulus parameters were delivered with a NIDAQ board controlled through Bonsai (https://open-ephys.org/). Animals were manually scored using Noldus EthoVision XT software (Noldus Information Technology), where grooming activity was plotted as the percentage of time spent grooming the antennae during red-light stimuli. The ablation of the antennae did not damage the rest of the head or lead to the mortality of the animals. Animals that died during the analysis window (7 - 14 dpa) were excluded.

### Automated antennal grooming behavior detection system

Frame-wise scoring of antennal grooming behavior was performed as an image-recognition task. Briefly, bouts of antennal grooming are a stereotypical pattern of foreleg movements directed toward the antennae that span several consecutive frames. This pattern was captured and compressed on a single image by clipping the grooming episode over time and computing the pixel-wise standard deviation through the stack of frames. Supervised training of an artificial neural network was used to recognize these compressed images belonging to one of two categories: grooming or no-grooming. An end-to-end system was established that takes a raw video as input and returns a frame-wise probability as output. Two neural networks and a set of image-processing functions wrapped in a single system were established, requiring minimal user input. The details of the system, together with the parameters used for training, validating, and testing are provided below.

### Network-1: Detection of the head and its fore region

To simplify the task of detecting grooming events, the region of interest was restricted to the forehead of the animal and the region in front of it. A deconvolutional network was trained to detect the head of the animal and to output the coordinates of its centroid.

#### Model

The deconvolutional network was developed in Python (V 3.7.4) using the PyTorch Library (V 1.13.0). The network is composed of six layers: 3 convolutional layers (channels: 32, 64, 128; kernel size: 3, 5, 3; max pooling: 2; stride: 1) and 6 deconvolutional layers (channels: 128, 64, 32; kernel size: 3, 5, 3; max unpooling: 2; stride: 1). We used the Kaiming weight initialization and ReLU activation functions as implemented in the Pytorch library.

#### Dataset

Raw videos are large (1280 × 1024 px) grayscale avi files. To reduce processing time, the network was trained, validated, and tested on lower resolution frames (320 × 256 px) extracted from the video files. 916 tuples (raw frame, binary mask of the head position) were used for training, 229 tuples for validation, and 250 tuples for testing the accuracy of the network.

#### Training, validation, and test

the network was trained to minimize the Mean Squared Error loss function with the ADAM optimizer on batches of 8 tuples for 14 epochs. The learning rate was fixed at 0.003. The coordinates of the binary mask’s centroid was used to probe the accuracy on the validation set. The Euclidean distance between the centroid of the binary mask and the max value of the confidence map produced by the network is used to evaluate the accuracy of the segmentation process. The distance (avgerage ± standard deviation) between centroids is 3.88 ± 3.48 pixels during validation and 4.33 ± 10.43 pixels in the test set. The minimum diameter of an animal’s head (e.g., side view relative to the camera) was estimated ~32 pixels. For this reason, a cut-off radius of 8 pixels (e.g., max distance between ground truth centroid and predicted centroid) was used to define the segmentation accuracy. According to this cut-off, the network correctly predicted the position of the head in 98.8 % of the test frames. The text file output of this network contains the frame-by-frame coordinates of the predicted position of the centroid of the head. In addition, the inference script automatically detects the onset/offset of the red-light stimuli by first taking the average pixel intensity of a 100 × 50 pixel region surrounding the LED on the initial 200 frames (light-off) and then computing the z-score for each frame in the video. Frames that are 2 z-scores above the average are defined as light-on frames. This procedure leads to a marginal underestimation (5.4 frames, mean absolute difference, data not shown) of the total number of light-on frames as compared to the measure taken by a human observer.

### Network-2: Grooming detection

The output of the deconvolutional network (e.g., the coordinates of the head’s centroid) was used to define an ROI centered on the head of the animal and its fore region. Following this crop, the pixel-wise standard deviation of 10 consecutive frames was computed, and the network classified the resulting image as either grooming or no-grooming. A sliding window (stride = 1 frame) was applied to the input to compute the frame-wise probability of grooming on the full length of the raw video. The network was initially trained on naïve animals. However, the inference on animals that underwent injury (e.g., antennal ablation) was suboptimal. Therefore, the network was trained on two distinct datasets (e.g., animals with or without antennae), leading to two distinct sets of weights that were independently deployed according to the experimental group (e.g., before or after injury). The animal condition did not affect the quality of the inference on head tracking performed by the deconvolutional network (Network-1).

#### Model

Pretrained ResNet50 (Pytorch: ResNet50_Weights.IMAGENET1K_V2) fine-tuned on our training datasets.

#### Dataset

Antennal grooming is defined as a stereotyped circular movement of the forelegs sweeping over the surface of the antennae. A trained experimenter manually labeled the grooming episodes using the manual scoring module of EthovisionXT16 (Noldus Information Technologies). This dataset represented the ground truth for training the ResNet50 and it required further processing before being used to train the network. First, the raw videos were processed with the deconvolutional network (see section ”Network-1: detection of the head and its fore region”) to extract the coordinates of the head’s centroid. Manual scoring of grooming events together with the coordinates of the head centroid were used to clip grooming bouts and crop frames (128 × 128 px) on a region centered on the head of the animal. For each grooming bout, the pixel-wise standard deviation was computed on a stack of 10 consecutive frames taken with a sliding window (stride = 1) over the bout. A sequence of 10 frames was used to capture short bouts (camera fps: 15 Hz, ~0.7 s) of grooming. The same process was repeated for frames labeled as “no_grooming”. These sets of images (e.g., grooming and no_grooming) were further inspected to remove images that could not be unambiguously attributed to one of the two categories. The images were then randomly allocated to one of three sets: train, validation, or test. Two distinct sets of images were obtained from the before and after-injury conditions. The before-injury set comprised 30 subjects (number of frames, train = 3499, validation = 943, test = 64), while the after-injury comprised 22 subjects (train = 1615, validation = 413, test = 495).

#### Training, validation, and test

The ResNet50 was trained for 41 epochs with a Cross Entropy loss function, the ADAM optimizer, and a decaying learning rate (gamma = 0.1, 7 epochs). The classification accuracy on the validation and test sets was 95.9 % and 93.5 %, respectively.

### Post-processing of grooming prediction

The ResNet50 model outputs a text file containing the framewise probability values for grooming behavior. However, because of the projection through time (10 consecutive frames) of the training set and the sliding window (*n* = 1), the probability value failed to capture sharp transitions between grooming states (e.g., onset-offset). For this reason, first, inference on the training set videos was run to get raw probability values; then these were binarized according to 7 probability threshold values θ ranging from 0.1 to 0.9 (Figure 3C, D) and for each frame with p > θ, the 3 consecutive frames were labeled as ‘grooming’. This procedure allowed to compensate for the bias introduced by the compression of 10 consecutive images without producing excess false positives. Finally, comparing the resulting grooming scores with manual scores allowed us to find an optimal probability threshold value that minimizes the difference between system and observer scores.

### Replication

Unless stated otherwise, at least 3 biological replications were performed for all experiments for each genotype and/or condition.

### Software and statistics

Image-J and Photoshop were used to process wing, *or22a^+^*, *JO^+^* and *orco^+^* images. The optogenetics section includes the software used for the analysis. The comparison between various observers vs. system, as well as the paired mean difference of automated antennal grooming detection, were plotted using https://www.estimationstats.com/#/. Unless otherwise stated, GraphPad Prism 9 was used for statistical analyses.

## Supplementary figure legend

**Supplementary figure S1.**
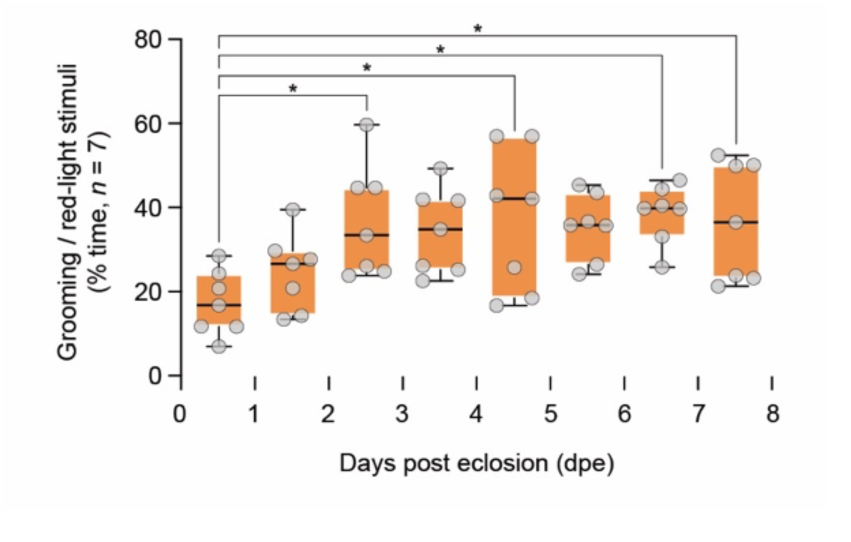
Time-course of optogenetically-evoked antennal grooming behavior in wild type after eclosion. Quantification of manually scored antennal grooming in wild-type flies with CsChrimson expressed in JO neurons between 0 to 7 days post eclosion (dpe). Data, % time of red-light stimuli (*n* = 7 animals). One-way ANOVA with Tukey’s multiple comparisons test. *, *p* < 0.05.

**Supplementary figure S2.**
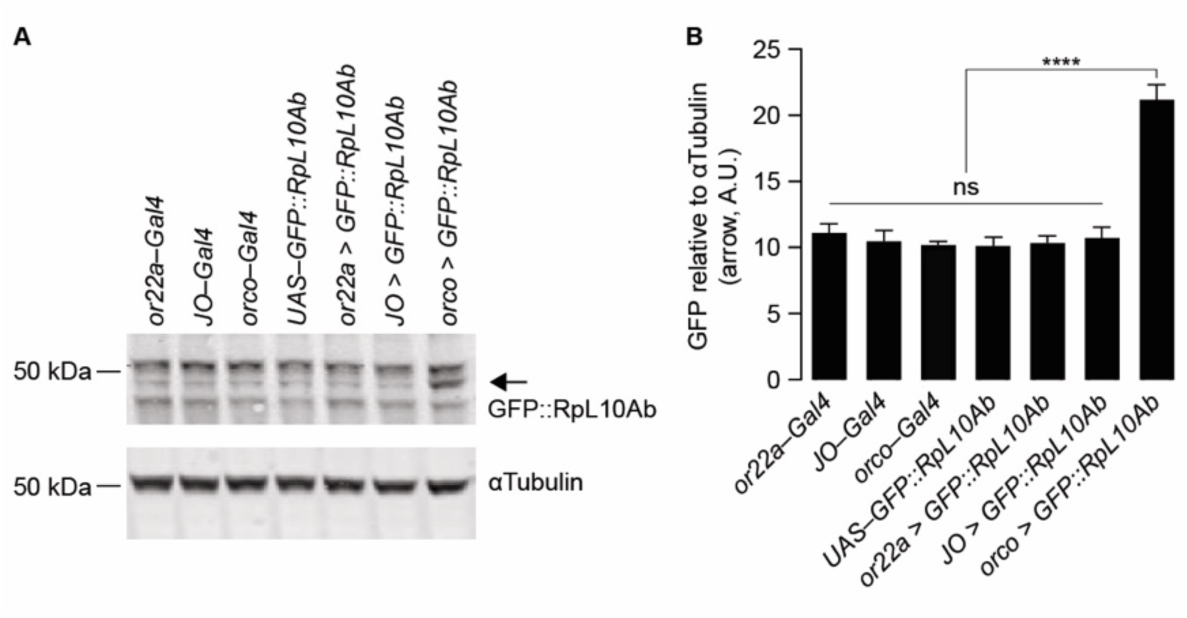
Olfactory organ projections as a large-scale resource to express and detect GFP-tagged ribosomal protein 10Ab. **A** Western blot of *Drosophila* heads with GFP-tagged ribosomal protein 10Ab (GFP::RpL10Ab) expressed in *orco^+^* neurons. 4 heads/lane; arrow, molecular weight of GFP::RpL10Ab. **B** Quantification of GFP immunoreactivity by densitometry. Data, mean ± SEM (*n* = 3, three replicates of three biological experiments). Arbitrary units, A.U.; One-way ANOVA with Tukey’s multiple comparisons test; ****, *p* < 0.0001; ns (not significant), *p* > 0.05.

**Supplementary figure S3.**
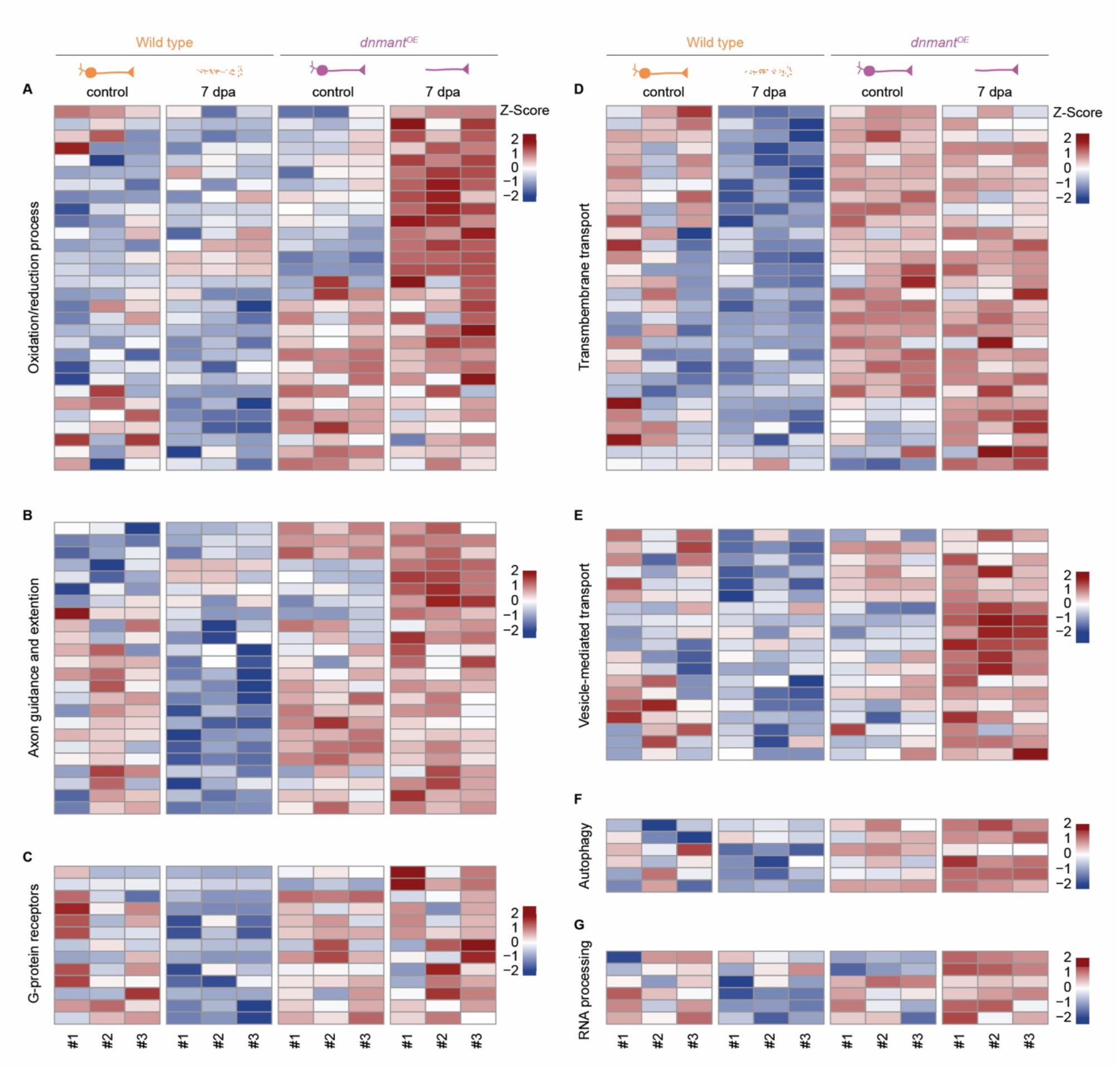
Heat maps of biological process GO terms significantly enriched in *dnmnat^OE^*7 dpa. **A** Heat map of oxidation/reduction process transcripts (*n =* 30). **B** Heat map of axon guidance and extension transcripts (*n =* 24). **C** Heat map of G-protein receptor transcripts (*n =* 13). **D** Heat map of transmembrane transport transcripts (*n =* 30). **E** Heat map of vesicle-mediated transport transcripts (*n =* 19). **F** Heat map of autophagy transcripts (*n =* 6). **G** Heat map of RNA processing transcripts (*n =* 6). Expression levels, Z-Score average.

**Supplementary figure S4.**
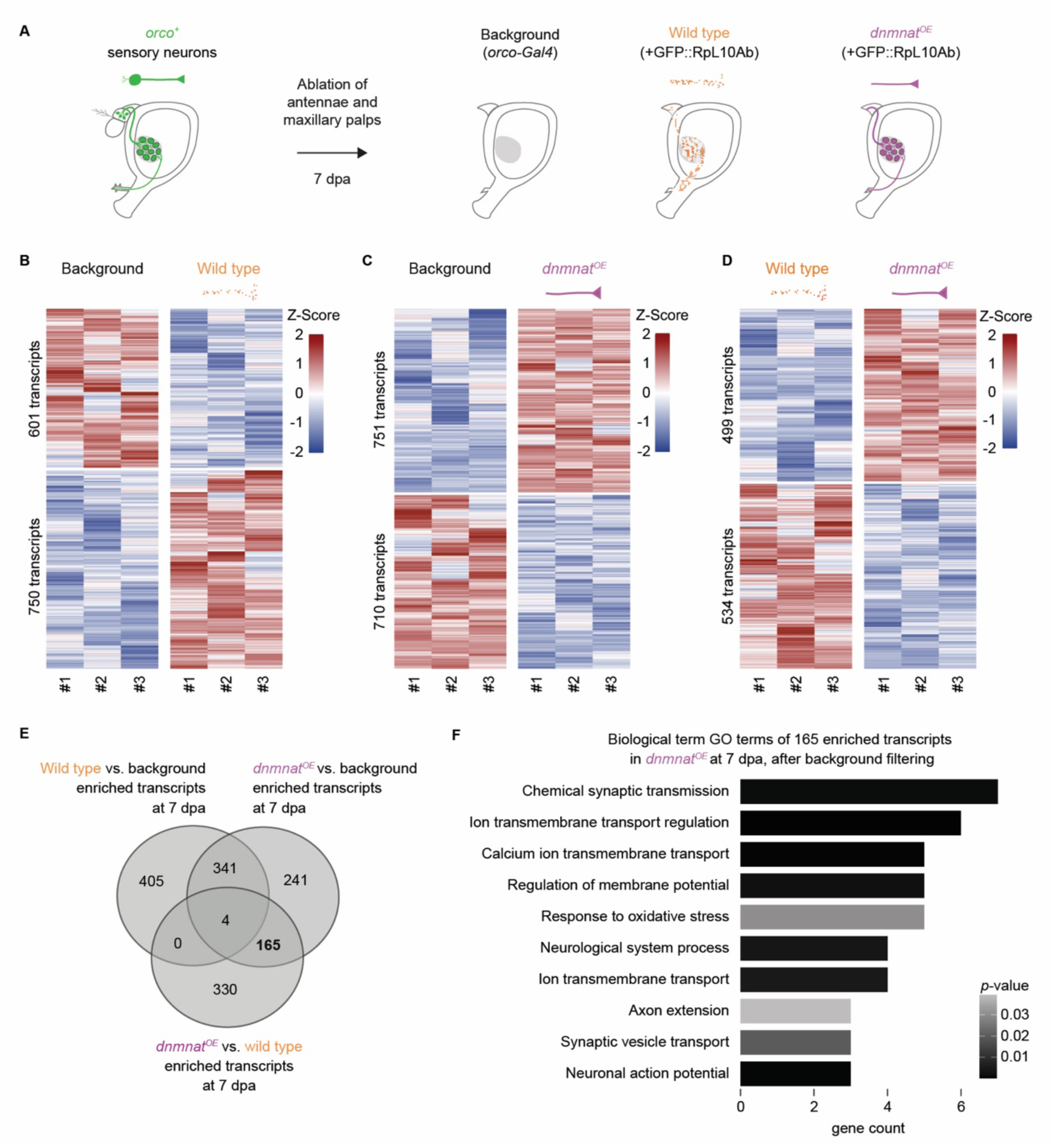
Background filtering of transcriptional profiles. **A** Strategy implemented for background filtering. **B** Heat map of 1351 transcripts differentially expressed in wild type (*orco > GFP::RpL10Ab*) compared to background (*orco–Gal4*) 7 dpa. **C** Heat map of 1461 transcripts differentially expressed in *dnmnat^OE^* (*orco > GFP::RpL10Ab*, *dnmnat*) compared to background (*orco–Gal4*) 7 dpa. **D** Heat map of 1033 transcripts differentially expressed in *dnmnat^OE^* (*orco > GFP::RpL10Ab*, *dnmnat*) compared to wild type (*orco > GFP::RpL10Ab*) 7 dpa. Data, Z-Score average. **E** Venn diagram of 750 transcripts enriched in wild type (compared to background), 751 transcripts enriched in *dnmnat^OE^* (compared to background), and 499 transcripts enriched in *dnmnat^OE^* (compared to wild type) 7 dpa. **F** Biological process GO term analysis of 165 enriched transcripts in *dnmnat^OE^* after background filtering (fold change value > 0.6); sorted and plotted by *p*-value.

**Supplementary figure S5.**
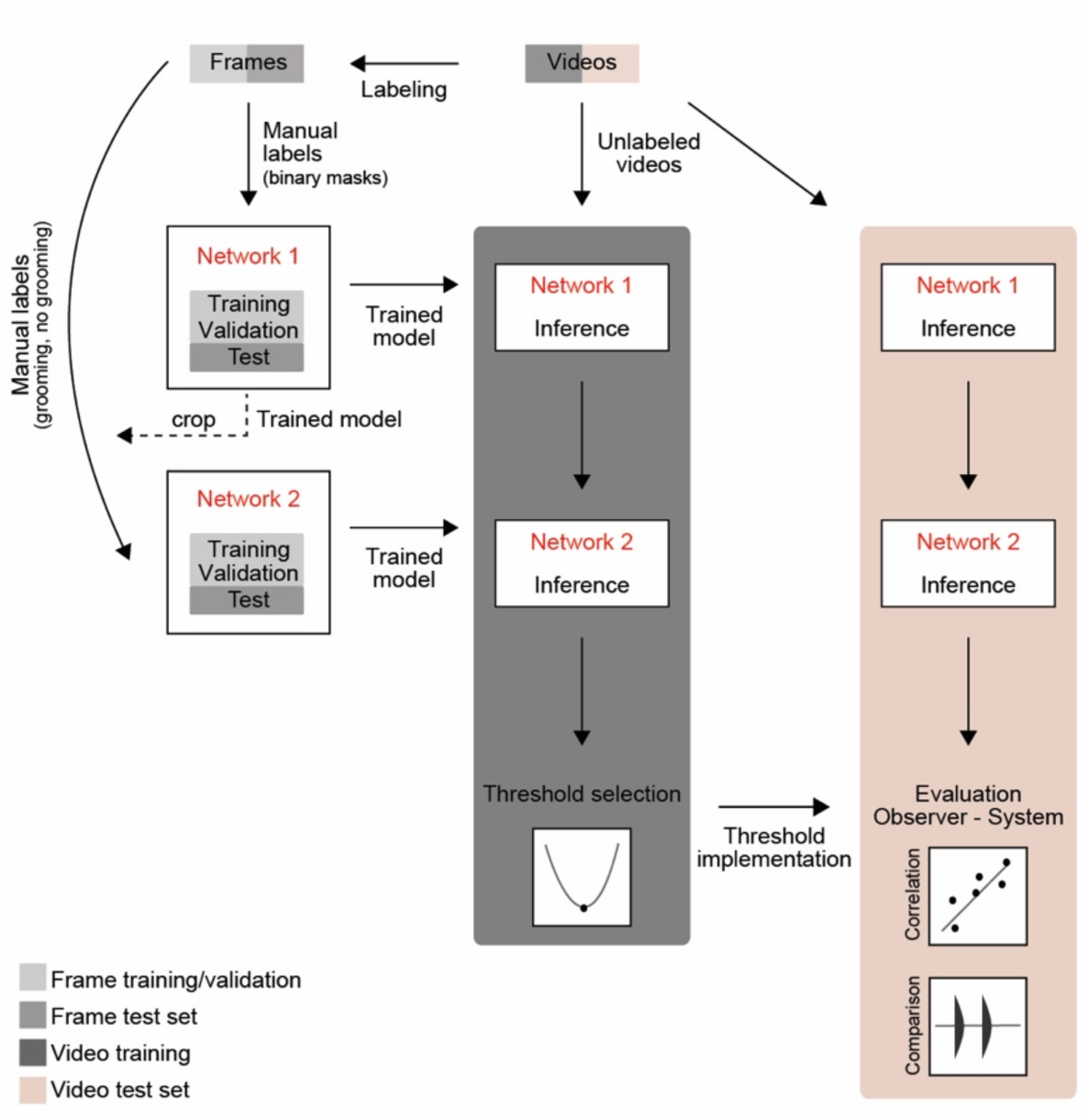
Illustration of Network 1 and 2 training and testing pipelines. Raw videos were split into training and test sets (grey and red, respectively). Video frames from the training set were manually labeled to identify the position of the head in the image (e.g. binary mask) and assign the grooming or no grooming labels. Network 1 was trained, validated, and tested on its own training set (frames). Network 2 was trained, validated, and tested on the labeled frames cropped by Network 1. Both networks, once trained, were used to determine the optimal threshold value of grooming detection using the same training set (raw videos). Finally, the test set (raw videos) was used to evaluate the accuracy of the system by comparing its performance with the scores of the observer.

**Supplementary figure S6.**
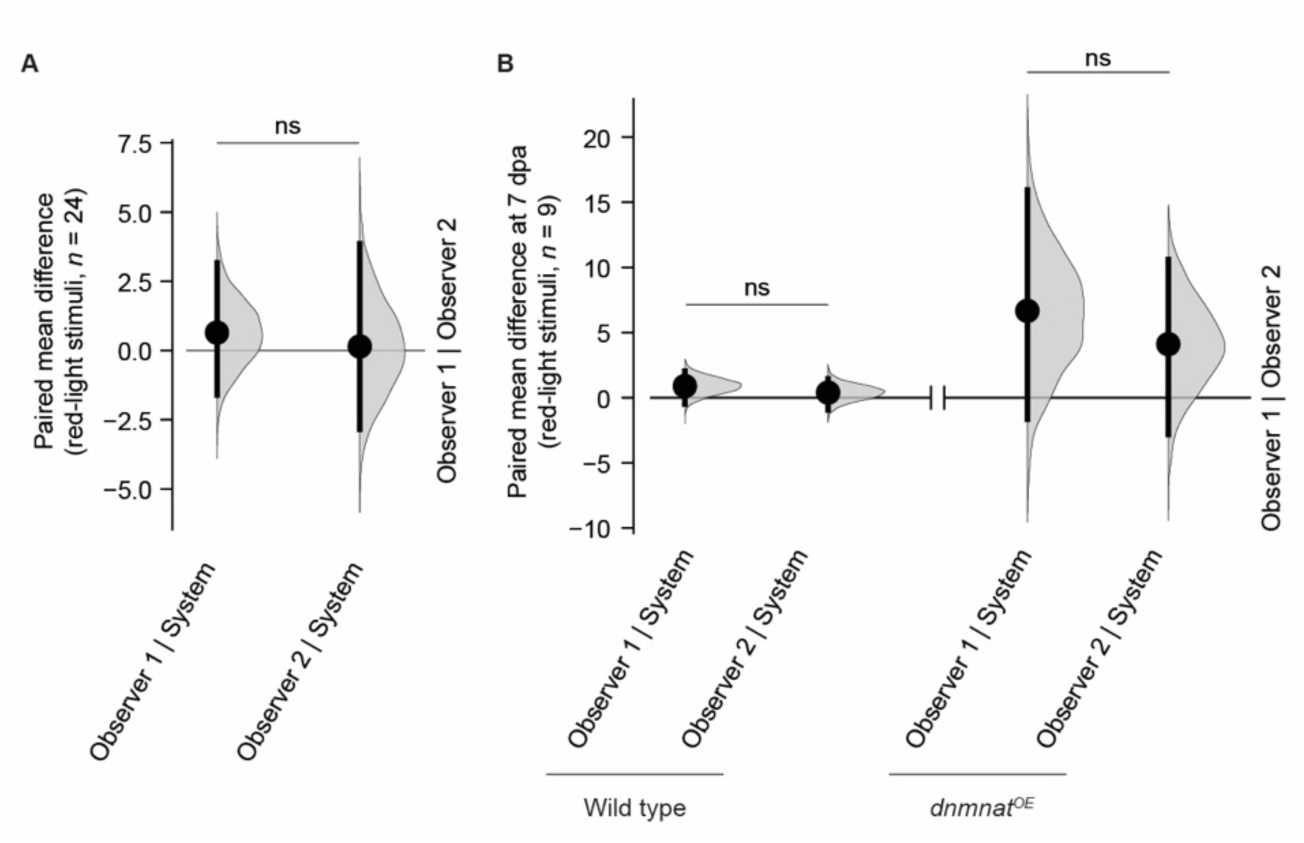
Scoring variability of antennal grooming between system and observers. **A** Standard deviation between grooming scores given by each observer to the same uninjured wild-type animal. **B** Standard deviation between grooming scores given by each observer to the same wild type and *dnmnat^OE^* at 7 dpa. The line break denotes the separation between the controls for each genotype. The plots illustrate the similarity between the standard deviations of observer 1 versus the system and observer 2 versus the system, as compared to the standard deviation observed among observers (horizontal black line, mean, number of frames). Non-parametric one-way ANOVA (Friedman test) with Dunn multiple comparisons test; ns (not significant), *p* > 0.05.

**Supplementary figure S7.**
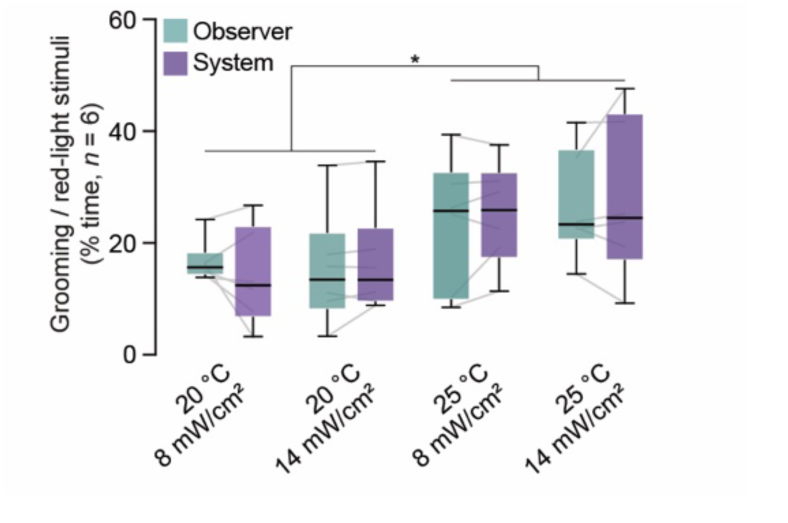
Detection of temperature-dependent changes in antennal grooming. Quantification of antennal grooming detection by observer and system in wild-type flies with CsChrimson expressed in JO neurons of 7-day-old animals (green and purple, respectively; probability, ≥ 0.3; % time of red-light stimuli; *n* = 6 animals). Three-way ANOVA; *, *p* < 0.05.

**Supplementary figure S8.**
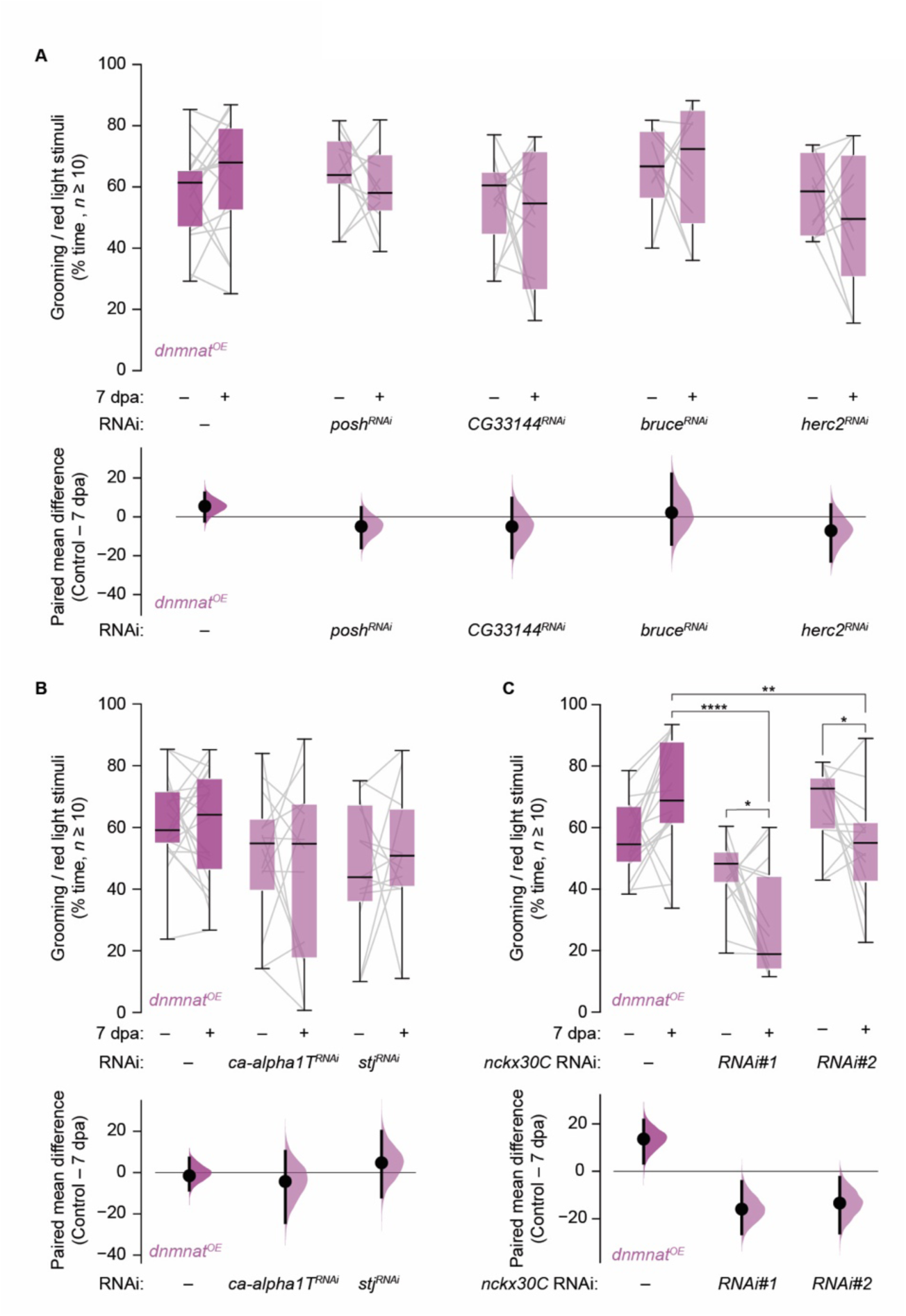
Protein ubiquitination and calcium transport candidate screen. **A** Top, protein ubiquitination candidate genes (Grooming as % time of red-light stimuli). Bottom, paired mean difference as bootstrap sampling distribution (Mean differences, dots; vertical error bars, 95% confidence intervals, respectively). **B** Top, calcium transport candidate genes; bottom, paired mean difference. **C** Top, validation of *nckx30C* by two different RNAi lines (#1 and #2, respectively); bottom paired mean difference. Paired two-tailed t-student and Two-Way ANOVA with Dunnett’s multiple comparisons test; ****, *p* < 0.0001; **, *p* < 0.01; *, *p* < 0.05.

## Video legend

**Video 1. Example of automated antennal grooming detection of a wild-type animal.** Left, recording of animal in arena exposed to red-light stimulus indicated by flashing diode (bottom). Square, network 1 head detection. Top right, network 2 antennal grooming detection (probability indicated in the upper left corner). Bottom right, probability plot. Wild-type animal expressing CsChrimson in JO neurons.

**Video 2. Example of automated antennal grooming detection of a wild-type animal after injury.** Left, recording of animal in arena exposed to red-light stimulus indicated by flashing diode (bottom). Square, network 1 head detection. Top right, network 2 antennal grooming detection (probability indicated in the upper left corner). Bottom right, probability plot. Wild-type animal expressing CsChrimson in JO neurons 7 dpa.

**Video 3. Example of automated antennal grooming detection of an animal expressing *dnmnat* in JO neurons after injury.** Left, recording of animal in arena exposed to red-light stimulus indicated by flashing diode (bottom). Square, network 1 head detection. Top right, network 2 antennal grooming detection (probability indicated in the upper left corner). Bottom right, probability plot. Wild-type animal expressing CsChrimson and *dnmnat* in JO neurons 7 dpa.

## Notes

### Competing Interest Statement

The authors have declared no competing interest.

### Summary of Updates

Revised abstract, and updated introduction, results, and discussion.

## Bibliography

Albuquerque, E. X., Deshpande, S. S., & Guth, L. (1978). Physiological properties of the innervated and denervated neuromuscular junction of hibernating and nonhibernating ground squirrels. Experimental Neurology, 62(2). 10.1016/0014-4886(78)90061-4

Alvarez, J. (2001). The autonomous axon: A model based on local synthesis of proteins. Biological Research, 34(2). 10.4067/S0716-97602001000200014

Anders, S., Pyl, P. T., & Huber, W. (2014). HTSeq—a Python framework to work with high-throughput sequencing data | Bioinformatics | Oxford Academic. Bioinformatics, 31(2).

Augustus Volney Waller. (1850). XX. Experiments on the section of the glossopharyngeal and hypoglossal nerves of the frog, and observations of the alterations produced thereby in the structure of their primitive fibres. Philosophical Transactions of the Royal Society of London, 140, 423–429. 10.1098/rstl.1850.0021

Bittner, G. D. (1988). Long term survival of severed distal axonal stumps in vertebrates and invertebrates. Integrative and Comparative Biology, 28(4). 10.1093/icb/28.4.1165

Bittner, G. D. (1991). Long-term survival of anucleate axons and its implications for nerve regeneration. In Trends in Neurosciences (Vol. 14, Issue 5). 10.1016/0166-2236(91)90104-3

Bolger, A. M., Lohse, M., & Usadel, B. (2014). Trimmomatic: A flexible trimmer for Illumina sequence data. Bioinformatics, 30(15). 10.1093/bioinformatics/btu170

Brazill, J. M., Li, C., Zhu, Y., & Zhai, R. G. (2017). NMNAT: It’s an NAD+ synthase… It’s a chaperone… It’s a neuroprotector. In Current Opinion in Genetics and Development (Vol. 44). 10.1016/j.gde.2017.03.014

Coleman, M. P., & Höke, A. (2020). Programmed axon degeneration: from mouse to mechanism to medicine. In Nature Reviews Neuroscience (Vol. 21, Issue 4). 10.1038/s41583-020-0269-3

Court, F. A., Hendriks, W. T. J., MacGillavry, H. D., Alvarez, J., & Van Minnen, J. (2008). Schwann cell to axon transfer of ribosomes: Toward a novel understanding of the role of glia in the nervous system. Journal of Neuroscience, 28(43). 10.1523/JNEUROSCI.2429-08.2008

Denny-Brown, D., Adams, R. D., Brenner, C., & Doherty, M. M. (1945). The pathology of injury to nerve induced by cold. Journal of Neuropathology and Experimental Neurology, 4(4). 10.1097/00005072-194504040-00001

Ding, Q., Cecarini, V., & Keller, J. N. (2007). Interplay between protein synthesis and degradation in the CNS: physiological and pathological implications. In Trends in Neurosciences (Vol. 30, Issue 1). 10.1016/j.tins.2006.11.003

Ederle, H., & Dormann, D. (2017). TDP-43 and FUS en route from the nucleus to the cytoplasm. In FEBS Letters (Vol. 591, Issue 11). 10.1002/1873-3468.12646

Essuman, K., Summers, D. W., Sasaki, Y., Mao, X., DiAntonio, A., & Milbrandt, J. (2017). The SARM1 Toll/Interleukin-1 Receptor Domain Possesses Intrinsic NAD+ Cleavage Activity that Promotes Pathological Axonal Degeneration. Neuron, 93(6). 10.1016/j.neuron.2017.02.022

Fang, Y., Soares, L., Teng, X., Geary, M., & Bonini, N. M. (2012). A novel drosophila model of nerve injury reveals an essential role of Nmnat in maintaining axonal integrity. Current Biology, 22(7). 10.1016/j.cub.2012.01.065

Gerdts, J., Brace, E. J., Sasaki, Y., DiAntonio, A., & Milbrandt, J. (2015). SARM1 activation triggers axon degeneration locally via NAD+ destruction. Science, 348(6233). 10.1126/science.1258366

Gilley, J., & Coleman, M. P. (2010). Endogenous Nmnat2 Is an Essential Survival Factor for Maintenance of Healthy Axons. PLoS Biology, 8(1). 10.1371/journal.pbio.1000300

Glock, C., Biever, A., Tushev, G., Nassim-Assir, B., Kao, A., Bartnik, I., Dieck, S. tom, & Schuman, E. M. (2021). The translatome of neuronal cell bodies, dendrites, and axons. Proceedings of the National Academy of Sciences of the United States of America, 118(43). 10.1073/pnas.2113929118

Hampel, S., Franconville, R., Simpson, J. H., & Seeds, A. M. (2015). A neural command circuit for grooming movement control. eLife, 4(September). 10.7554/eLife.08758

Hampel, S., McKellar, C. E., Simpson, J. H., & Seeds, A. M. (2017). Simultaneous activation of parallel sensory pathways promotes a grooming sequence in drosophila. eLife, 6. 10.7554/eLife.28804

Holt, C. E., Martin, K. C., & Schuman, E. M. (2019). Local translation in neurons: visualization and function. In Nature Structural and Molecular Biology (Vol. 26, Issue 7). 10.1038/s41594-019-0263-5

Hu, Y., Flockhart, I., Vinayagam, A., Bergwitz, C., Berger, B., Perrimon, N., & Mohr, S. E. (2011). An integrative approach to ortholog prediction for disease-focused and other functional studies. BMC Bioinformatics, 12. 10.1186/1471-2105-12-357

Huang, D. W., Sherman, B. T., & Lempicki, R. A. (2009). Systematic and integrative analysis of large gene lists using DAVID bioinformatics resources. Nature Protocols, 4(1). 10.1038/nprot.2008.211

Ishiguro, A., Kimura, N., Watanabe, Y., Watanabe, S., & Ishihama, A. (2016). TDP-43 binds and transports G-quadruplex-containing mRNAs into neurites for local translation. Genes to Cells, 21(5). 10.1111/gtc.12352

Iwata, A., Stys, P. K., Wolf, J. A., Chen, X. H., Taylor, A. G., Meaney, D. F., & Smith, D. H. (2004). Traumatic axonal injury induces proteolytic cleavage of the voltage-gated sodium channels modulated by tetrodotoxin and protease inhibitors. Journal of Neuroscience, 24(19). 10.1523/JNEUROSCI.0515-03.2004

Jarome, T. J., & Helmstetter, F. J. (2014). Protein degradation and protein synthesis in long-term memory formation. In Frontiers in Molecular Neuroscience (Vol. 7, Issue JUNE). 10.3389/fnmol.2014.00061

Jung, H., Yoon, B. C., & Holt, C. E. (2012). Axonal mRNA localization and local protein synthesis in nervous system assembly, maintenance and repair. In Nature Reviews Neuroscience (Vol. 13, Issue 5). 10.1038/nrn3210

Jung, J., Ohk, J., Kim, H., Holt, C. E., Park, H. J., & Jung, H. (2023). mRNA transport, translation, and decay in adult mammalian central nervous system axons. Neuron, 111(5), 650–668.e4. 10.1016/j.neuron.2022.12.015

Kim, D., Langmead, B., & Salzberg, S. L. (2015). HISAT: A fast spliced aligner with low memory requirements. Nature Methods, 12(4). 10.1038/nmeth.3317

Larsson, M. C., Domingos, A. I., Jones, W. D., Chiappe, M. E., Amrein, H., & Vosshall, L. B. (2004). Or83b encodes a broadly expressed odorant receptor essential for Drosophila olfaction. Neuron, 43(5). 10.1016/j.neuron.2004.08.019

Lee, T., & Luo, L. (1999). Mosaic analysis with a repressible neurotechnique cell marker for studies of gene function in neuronal morphogenesis. Neuron, 22(3). 10.1016/S0896-6273(00)80701-1

Llobet Rosell, A., & Neukomm, L. J. (2019). Axon death signalling in Wallerian degeneration among species and in disease. In Open Biology (Vol. 9, Issue 8). 10.1098/rsob.190118

Llobet Rosell, A., Paglione, M., Gilley, J., Kocia, M., Perillo, G., Gasparrini, M., Cialabrini, L., Raffaelli, N., Angeletti, C., Orsomando, G., Wu, P. H., Coleman, M. P., Loreto, A., & Neukomm, L. J. (2022). The NAD+ precursor NMN activates dSarm to trigger axon degeneration in Drosophila. eLife, 11. 10.7554/ELIFE.80245

Love, M. I., Huber, W., & Anders, S. (2014). Moderated estimation of fold change and dispersion for RNA-seq data with DESeq2. Genome Biology, 15(12). 10.1186/s13059-014-0550-8

Lunn, E. R., Perry, V. H., Brown, M. C., Rosen, H., & Gordon, S. (1989). Absence of Wallerian Degeneration does not Hinder Regeneration in Peripheral Nerve. European Journal of Neuroscience, 1(1). 10.1111/j.1460-9568.1989.tb00771.x

MacDonald, J. M., Beach, M. G., Porpiglia, E., Sheehan, A. E., Watts, R. J., & Freeman, M. R. (2006). The Drosophila Cell Corpse Engulfment Receptor Draper Mediates Glial Clearance of Severed Axons. Neuron, 50(6). 10.1016/j.neuron.2006.04.028

Mack, T. G. A., Reiner, M., Beirowski, B., Mi, W., Emanuelli, M., Wagner, D., Thomson, D., Gillingwater, T., Court, F., Conforti, L., Fernando, F. S., Tarlton, A., Andressen, C., Addicks, K., Magni, G., Ribchester, R. R., Perry, V. H., & Coleman, M. P. (2001). Wallerian degeneration of injured axons and synapses is delayed by a Ube4b/Nmnat chimeric gene. Nature Neuroscience, 4(12). 10.1038/nn770

Neukomm, L. J., Burdett, T. C., Gonzalez, M. A., Züchner, S., & Freeman, M. R. (2014). Rapid in vivo forward genetic approach for identifying axon death genes in Drosophila. Proceedings of the National Academy of Sciences of the United States of America, 111(27). 10.1073/pnas.1406230111

Neukomm, L. J., Burdett, T. C., Seeds, A. M., Hampel, S., Coutinho-Budd, J. C., Farley, J. E., Wong, J., Karadeniz, Y. B., Osterloh, J. M., Sheehan, A. E., & Freeman, M. R. (2017). Axon Death Pathways Converge on Axundead to Promote Functional and Structural Axon Disassembly. Neuron, 95(1). 10.1016/j.neuron.2017.06.031

Osterloh, J. M., Yang, J., Rooney, T. M., Fox, A. N., Adalbert, R., Powell, E. H., Sheehan, A. E., Avery, M. A., Hackett, R., Logan, M. A., MacDonald, J. M., Ziegenfuss, J. S., Milde, S., Hou, Y. J., Nathan, C., Ding, A., Brown, R. H., Conforti, L., Coleman, M., … Freeman, M. R. (2012). dSarm/Sarm1 is required for activation of an injury-induced axon death pathway. Science, 337(6093). 10.1126/science.1223899

Ostroff, L. E., Santini, E., Sears, R., Deane, Z., Kanadia, R. N., Ledoux, J. E., Lhakhang, T., Tsirigos, A., Heguy, A., & Klann, E. (2019). Axon TRAP reveals learning-associated alterations in cortical axonal mRNAs in the lateral amgydala. eLife, 8. 10.7554/eLife.51607

Paglione, M., Llobet Rosell, A., Chatton, J. Y., & Neukomm, L. J. (2020a). Morphological and Functional Evaluation of Axons and their Synapses during Axon Death in Drosophila melanogaster. J. Vis. Exp, 157, 60865. 10.3791/60865

Paglione, M., Llobet Rosell, A., Chatton, J.-Y., & Neukomm, L. J. (2020b). Morphological and Functional Evaluation of Axons and their Synapses during Axon Death in <EM>Drosophila melanogaster</EM>. Journal of Visualized Experiments, 157. 10.3791/60865-v

Raiders, S., Han, T., Scott-Hewitt, N., Kucenas, S., Lew, D., Logan, M. A., & Singhvi, A. (2021). Engulfed by glia: Glial pruning in development, function, and injury across species. Journal of Neuroscience, 41(5). 10.1523/JNEUROSCI.1660-20.2020

Rishal, I., & Fainzilber, M. (2014). Axon-soma communication in neuronal injury. In Nature Reviews Neuroscience (Vol. 15, Issue 1). 10.1038/nrn3609

Rozenbaum, M., Rajman, M., Rishal, I., Koppel, I., Koley, S., Medzihradszky, K. F., Oses-Prieto, J. A., Kawaguchi, R., Amieux, P. S., Burlingame, A. L., Coppola, G., & Fainzilber, M. (2018). Translatome regulation in neuronal injury and axon regrowth. eNeuro, 5(2). 10.1523/ENEURO.0276-17.2018

Sapar, M. L., & Han, C. (2019). Die in pieces: How Drosophila sheds light on neurite degeneration and clearance. In Journal of Genetics and Genomics (Vol. 46, Issue 4). 10.1016/j.jgg.2019.03.010

Schaumburg, H., Byck, R., Herman, R., & Rosengart, C. (1967). Peripheral Nerve Damage by Cold. Archives of Neurology, 16(1). 10.1001/archneur.1967.00470190107013

Schnell, M. J., McGettigan, J. P., Wirblich, C., & Papaneri, A. (2010). The cell biology of rabies virus: Using stealth to reach the brain. In Nature Reviews Microbiology (Vol. 8, Issue 1). 10.1038/nrmicro2260

Stirling, D. P., & Stys, P. K. (2010). Mechanisms of axonal injury: internodal nanocomplexes and calcium deregulation. In Trends in Molecular Medicine (Vol. 16, Issue 4). 10.1016/j.molmed.2010.02.002

Sundaramoorthy, V., Green, D., Locke, K., O’Brien, C. M., Dearnley, M., & Bingham, J. (2020). Novel role of SARM1 mediated axonal degeneration in the pathogenesis of rabies. PLoS Pathogens, 16(2). 10.1371/journal.ppat.1008343

Terenzio, M., Koley, S., Samra, N., Rishal, I., Zhao, Q., Sahoo, P. K., Urisman, A., Marvaldi, L., Oses-Prieto, J. A., Forester, C., Gomes, C., Kalinski, A. L., Di Pizio, A., Doron-Mandel, E., Perry, R. B. T., Koppel, I., Twiss, J. L., Burlingame, A. L., & Fainzilber, M. (2018). Locally translated mTOR controls axonal local translation in nerve injury. Science, 359(6382). 10.1126/science.aan1053

Thelen, M. P., & Kye, M. J. (2020). The Role of RNA Binding Proteins for Local mRNA Translation: Implications in Neurological Disorders. In Frontiers in Molecular Biosciences (Vol. 6). 10.3389/fmolb.2019.00161

Thomas, A., Lee, P. J., Dalton, J. E., Nomie, K. J., Stoica, L., Costa-Mattioli, M., Chang, P., Nuzhdin, S., Arbeitman, M. N., & Dierick, H. A. (2012). A versatile method for cell-specific profiling of translated mrnas in Drosophila. PLoS ONE, 7(7). 10.1371/journal.pone.0040276

Vosshall, L. B., Wong, A. M., & Axel, R. (2000). An olfactory sensory map in the fly brain. Cell, 102(2). 10.1016/S0092-8674(00)00021-0

Wang, J. W., Wong, A. M., Flores, J., Vosshall, L. B., & Axel, R. (2003). Two-photon calcium imaging reveals an odor-evoked map of activity in the fly brain. Cell, 112(2). 10.1016/S0092-8674(03)00004-7

Xiong, X., Hao, Y., Sun, K., Li, J., Li, X., Mishra, B., Soppina, P., Wu, C., Hume, R. I., & Collins, C. A. (2012). The Highwire Ubiquitin Ligase Promotes Axonal Degeneration by Tuning Levels of Nmnat Protein. PLoS Biology, 10(12). 10.1371/journal.pbio.1001440

Yamaguchi, M., Lee, I. soon, Jantrapirom, S., Suda, K., & Yoshida, H. (2021). Drosophila models to study causative genes for human rare intractable neurological diseases. In Experimental Cell Research (Vol. 403, Issue 1). 10.1016/j.yexcr.2021.112584

Yamaguchi, M., & Takashima, H. (2018). Drosophila charcot-marie-tooth disease models. In Advances in Experimental Medicine and Biology (Vol. 1076). 10.1007/978-981-13-0529-0_7

Yang, J., Wu, Z., Renier, N., Simon, D. J., Uryu, K., Park, D. S., Greer, P. A., Tournier, C., Davis, R. J., & Tessier-Lavigne, M. (2015). Pathological axonal death through a Mapk cascade that triggers a local energy deficit. Cell, 160(1–2). 10.1016/j.cell.2014.11.053

Yoon, B. C., Jung, H., Dwivedy, A., O’Hare, C. M., Zivraj, K. H., & Holt, C. E. (2012). Local translation of extranuclear lamin B promotes axon maintenance. Cell, 148(4). 10.1016/j.cell.2011.11.064

Zhai, R. G., Cao, Y., Hiesinger, P. R., Zhou, Y., Mehta, S. Q., Schulze, K. L., Verstreken, P., & Bellen, H. J. (2006). Drosophila NMNAT maintains neural integrity independent of its NAD synthesis activity. PLoS Biology, 4(12). 10.1371/journal.pbio.0040416

Zhai, R. G., Rizzi, M., & Garavaglia, S. (2009). Nicotinamide/nicotinic acid mononucleotide adenylyltransferase, new insights into an ancient enzyme. In Cellular and Molecular Life Sciences (Vol. 66, Issue 17). 10.1007/s00018-009-0047-x

Zhang, K., Coyne, A. N., & Lloyd, T. E. (2018). Drosophila models of amyotrophic lateral sclerosis with defects in RNA metabolism. In Brain Research (Vol. 1693). 10.1016/j.brainres.2018.04.043

ZhenXian Niou, S. Y. A. S. H. R. V. O. P. G. M. P. C. V. O. P.-C. L. (2022). NMNAT2 in cortical glutamatergic neurons exerts both cell and non-cell autonomous influences to shape cortical development and to maintain neuronal health. bioRxiv.

Zhulyn, O., Rosenblatt, H. D., Shokat, L., Dai, S., Kuzuoglu-Öztürk, D., Zhang, Z., Ruggero, D., Shokat, K. M., & Barna, M. (2023). Evolutionarily divergent mTOR remodels translatome for tissue regeneration. Nature, 620(7972), 163–171. 10.1038/s41586-023-06365-1

