## Supplementary Figures for "Local translation is engaged to sustain synaptic function in impaired Wallerian degeneration"

Paglione *et al.*, Supplementary figure S1

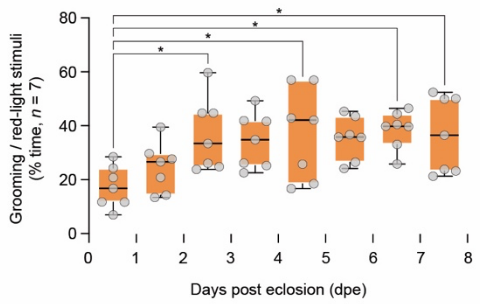

Paglione *et al.*, Supplementary figure S2

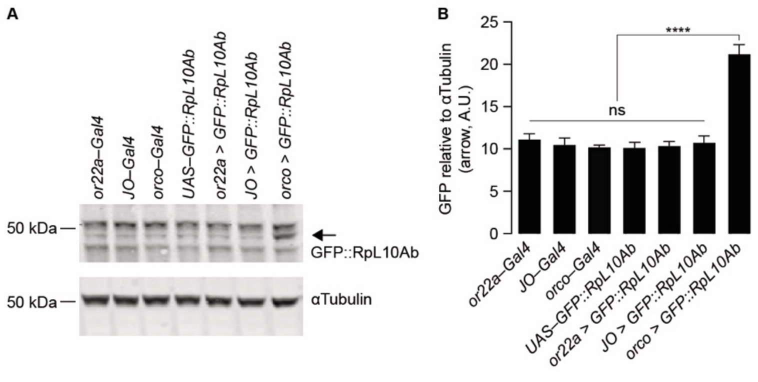

Paglione *et al.*, Supplementary figure S3

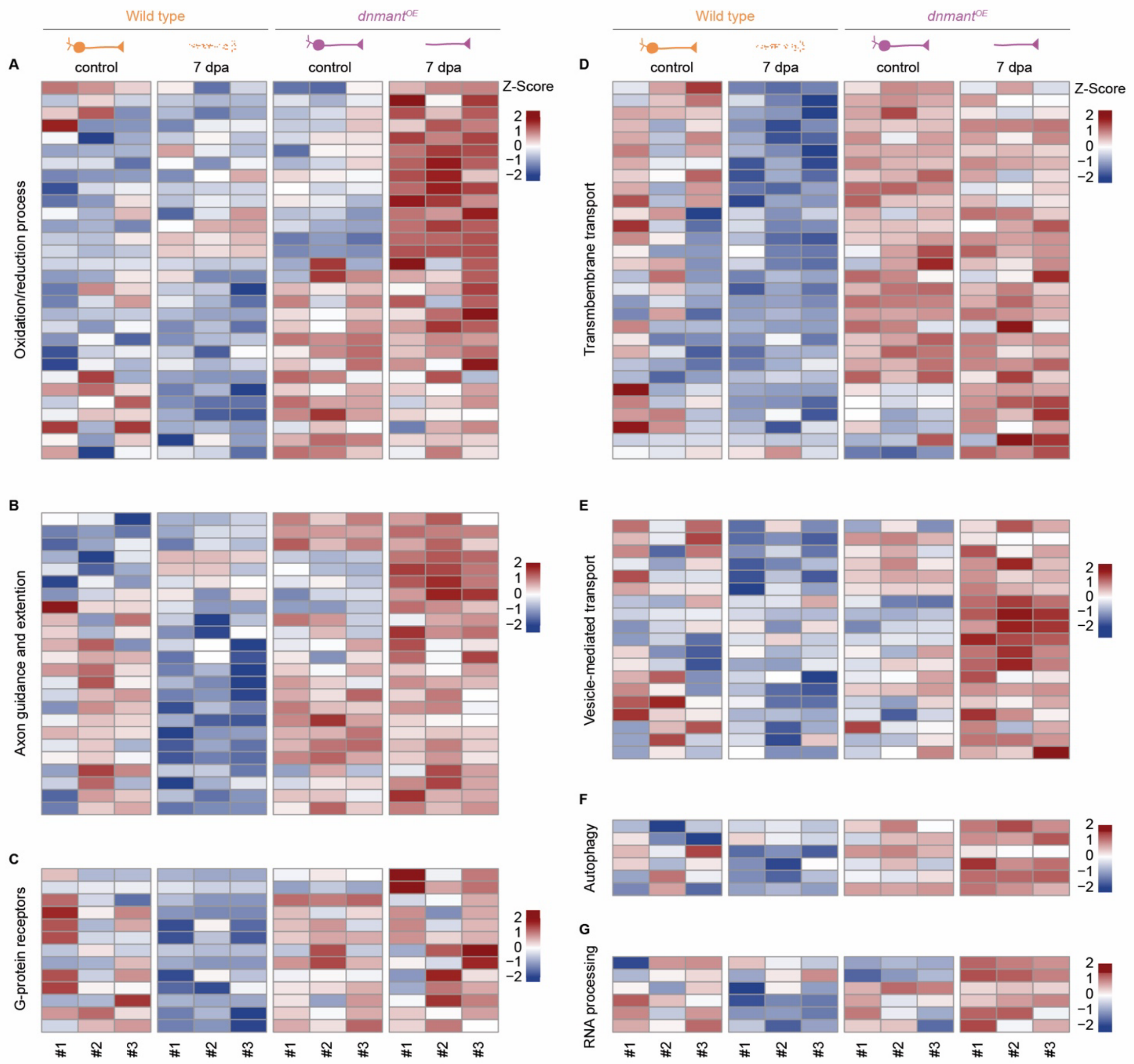

Paglione *et al.*, Supplementary figure S4

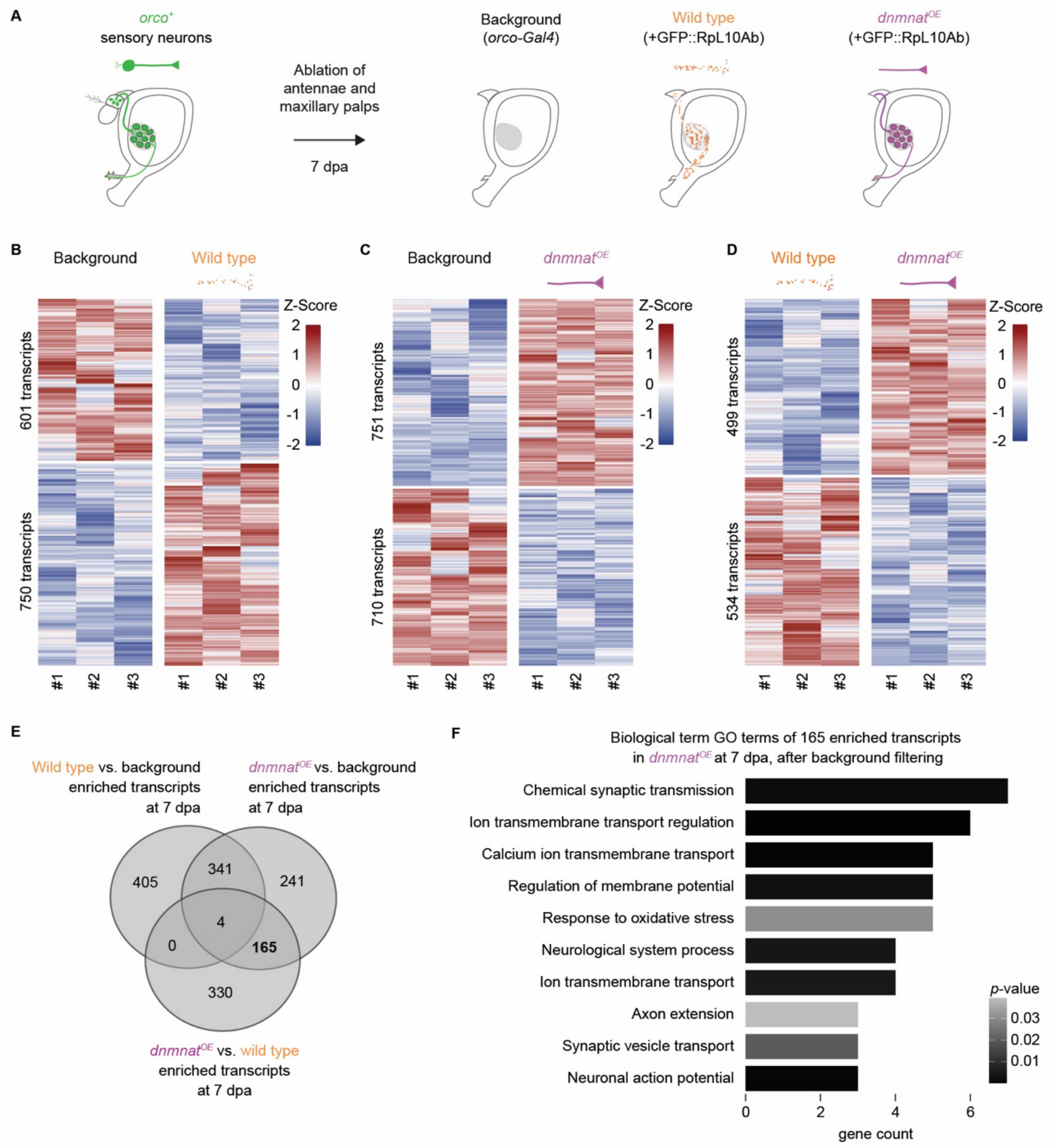

Paglione *et al.*, Supplementary figure S5

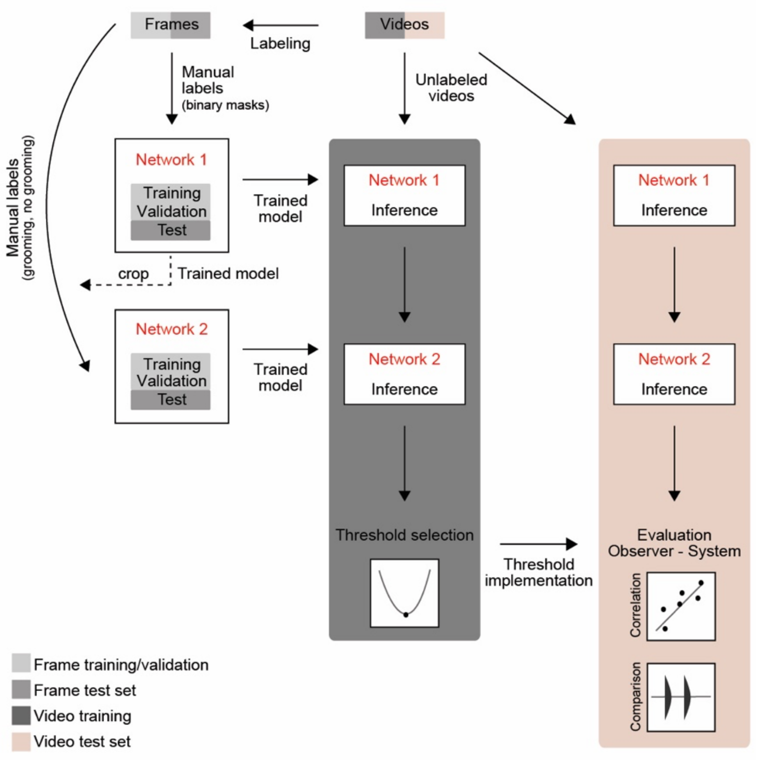

Paglione *et al.*, Supplementary figure S6

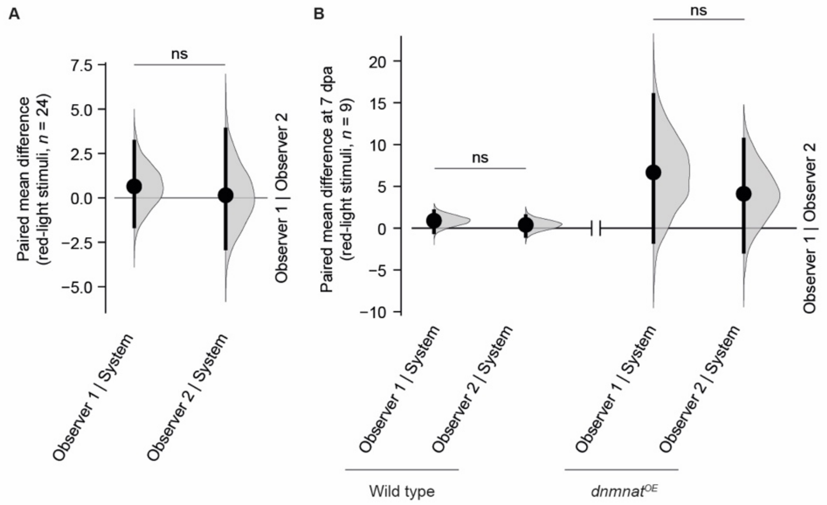

Paglione *et al.*, Supplementary figure S7

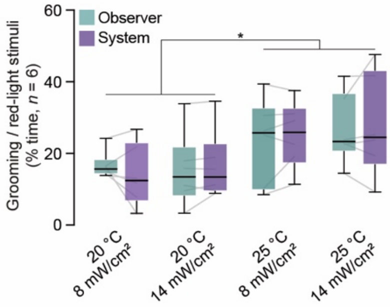

Paglione *et al.*, Supplementary figure S8

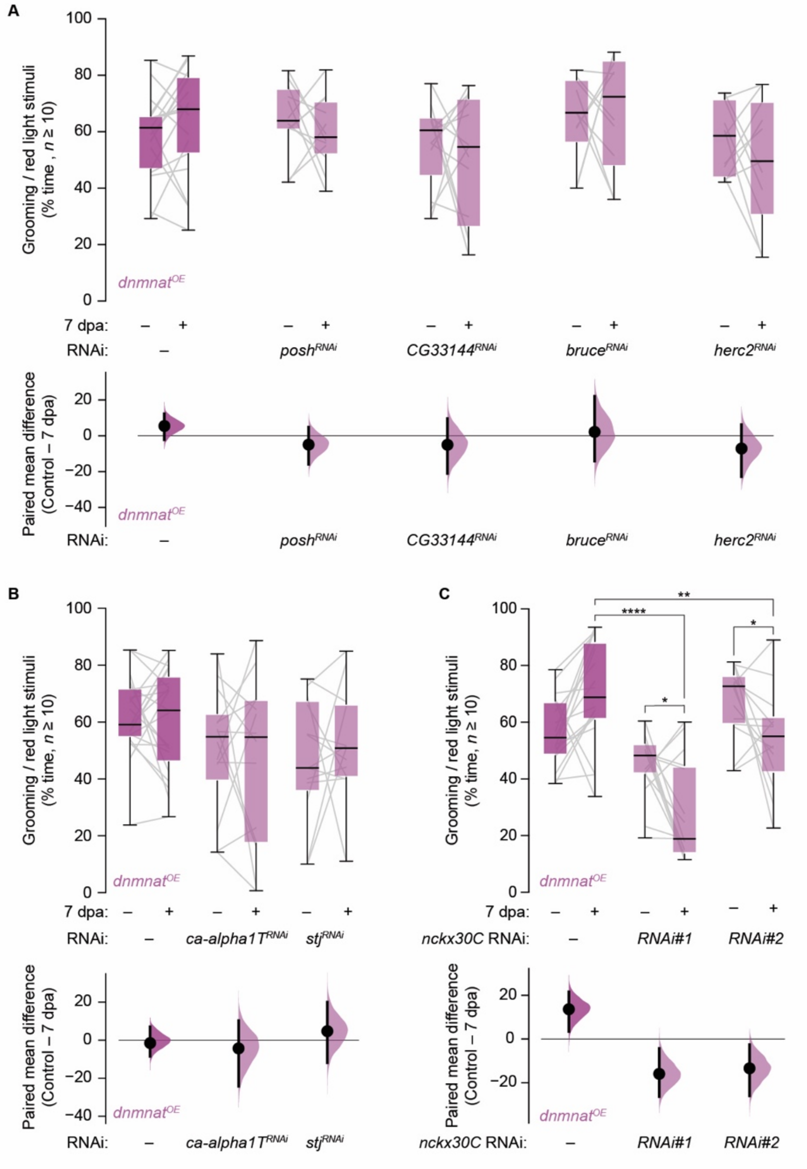
