## Supplementary material for "Local translation is engaged to sustain synaptic function in impaired Wallerian degeneration": Genotype list

### Figure 1.

**A-B:** Wild type: *w ; dpr1,mCD8::GFP,aseFLP*$^{2b}$*/+ ; FRT2A/tub-Gal80,FRT2A*
**A-B:** *dnmnat*$^{OE}$: *w ; dpr1,mCD8::GFP,aseFLP*$^{2b}$*/5xUAS–dnmnat ; FRT2A/tub-Gal80,FRT2A*
**C-D:** Wild type: *w ; or22a–Gal4,5xUAS–mCD8::GFP/+ ;*
**C-D:** *dnmnat*$^{OE}$: *w ; or22a–Gal4,5xUAS–mCD8::GFP/5xUAS–dnmnat ;*
**E-F:** Wild type: *w ; 5xUAS–mCD8::GFP/+ ; GMR60E02–Gal4 /+ ;*
**E-F:** *dnmnat*$^{OE}$: *w ; 5xUAS–mCD8::GFP/5xUAS–dnmnat ; GMR60E02–Gal4/+ ;*
**G:** Wild type: *w ; 20xUAS–IVS–CsChrimson::mVenus/+ ; GMR60E02–Gal4/+ ;*
**G:** *dnmnat*$^{OE}$: *w ; 20xUAS–IVS–CsChrimson::mVenus/5xUAS–dnmnat ; GMR60E02–Gal4/+ ;*
**H-I:** Wild type: *w ; orco–Gal4/+ ; 5xUAS–mCD8::GFP/+ ;*
**H-I:** *dnmnat*$^{OE}$: *w ; orco–Gal4/5xUAS–dnmnat ; 5xUAS–mCD8::GFP/+ ;*

### Figure 2.

**A-F:** Wild type: *w ; orco–Gal4/+ ; 5xUAS–GFP::RpL10Ab/+ ;*
**A-F:** *dnmnat*$^{OE}$: *w ; orco–Gal4/5xUAS–dnmnat ; 5xUAS–GFP::RpL10Ab/+ ;*

### Figure 3.

**A-H:** Wild type: *w ; 20xUAS–IVS–CsChrimson::mVenus/+ ; GMR60E02–Gal4/+ ;*
**F-H:** *dnmnat*$^{OE}$: *w ; 20xUAS-IVS-CsChrimson::mVenus/5xUAS–dnmnat ; GMR60E02–Gal4/+ ;*

### Figure 4.

**A-D:** Control: *w ; 20xUAS–IVS–CsChrimson::mVenus;5xUAS–dnmnat/+ ; GMR60E02–Gal4/+ ;*
**A-D:** RNAi(II): *w ; 20xUAS–IVS–CsChrimson::mVenus;5xUAS–dnmnat/5xUAS–RNAi ; GMR60E02–Gal4/+ ;*
**A-D:** RNAi(III): *w ; 20xUAS–IVS–CsChrimson::mVenus;5xUAS–dnmnat/+ ; GMR60E02–Gal4/5xUAS–RNAi ;*

### Figure 5.

**B:** Control: *w ; 20xUAS–IVS–CsChrimson::mVenus;5xUAS–dnmnat/+ ; GMR60E02–Gal4/+ ;*
**B:** RNAi(II): *w ; 20xUAS–IVS–CsChrimson::mVenus;5xUAS–dnmnat/5xUAS–RNAi ; GMR60E02–Gal4/+ ;*
**B:** RNAi(III): *w ; 20xUAS–IVS–CsChrimson::mVenus;5xUAS–dnmnat/+ ; GMR60E02–Gal4/5xUAS–RNAi ;*

### Supplementary figure 1.

Wild type: *w ; 20xUAS–IVS–CsChrimson::mVenus/+ ; GMR60E02–Gal4/+ ;*

### Supplementary figure 2.

**A-B:** *or22a–Gal4*: *w ; or22a–Gal4/+ ;*
**A-B:** *GMR60E02–Gal4:* *w ; ; GMR60E02–Gal4/+ ;*
**A-B:** *orco–Gal4*: *w ; orco–Gal4/+ ;*
**A-B:** *UAS–GFP::RpL10Ab*: *w ; ; 5xUAS–GFP::RpL10Ab/+ ;*
**A-B:** *or22a > GFP::RpL10Ab*: *w ; or22a–Gal4/+ ; 5xUAS–GFP::RpL10Ab/+ ;*
**A-B:** *GMR60E02 > GFP::RpL10Ab*: *w ; ; GMR60E02–Gal4/5xUAS–GFP::RpL10Ab ;*
**A-B:** *orco > GFP::RpL10Ab*: *w ; orco–Gal4/+ ; 5xUAS–GFP::RpL10Ab/+ ;*

### Supplementary figure 3.

**A-G:** Wild type: *w ; orco–Gal4/+ ; 5xUAS–GFP::RpL10Ab/+ ;*
**A-G:** *dnmnat*$^{OE}$: *w ; orco–Gal4/5xUAS–dnmnat ; 5xUAS–GFP::RpL10Ab/+ ;*

### Supplementary figure 4.

**A-F:** background: *w ; orco–Gal4/+ ; ;*
**A-F:** Wild type: *w ; orco–Gal4/+ ; 5xUAS–GFP::RpL10Ab/+ ;*
**A-F:** *dnmnat*$^{OE}$: *w ; orco–Gal4/5xUAS–dnmnat ; 5xUAS–GFP::RpL10Ab/+ ;*

### Supplementary figure 6.

**A** Wild type: *w ; 20xUAS–IVS–CsChrimson::mVenus/+ ; GMR60E02–Gal4/+ ;*
**B:** Wild type: *w ; 20xUAS–IVS–CsChrimson::mVenus/+ ; GMR60E02–Gal4/+ ;*
**B:** *dnmnat*$^{OE}$: *w ; 20xUAS–IVS–CsChrimson::mVenus/5xUAS–dnmnat ; GMR60E02–Gal4/+ ;*

### Supplementary figure 7.

Wild type: *w ; 20xUAS–IVS–CsChrimson::mVenus/+ ; GMR60E02–Gal4/+ ;*

### Supplementary figure 8.

**A-C:** Control: *w,20xUAS–IVS–CsChrimson::mVenus ; 5xUAS–dnmnat/+ ; GMR60E02–Gal4/+ ;*
**A-C:** RNAi(II): *w,20xUAS–IVS–CsChrimson::mVenus ; 5xUAS–dnmnat/5xUAS-RNAi ; GMR60E02–Gal4/+ ;*
**A-C:** RNAi(III): *w,20xUAS–IVS–CsChrimson::mVenus ; 5xUAS–dnmnat/+ ; GMR60E02–Gal4/5xUAS-RNAi ;*

### Video 1.

Wild type: *w ; 20xUAS–IVS–CsChrimson::mVenus/+ ; GMR60E02–Gal4/+ ;*

### Video 2.

Wild type: *w ; 20xUAS–IVS–CsChrimson::mVenus/+ ; GMR60E02–Gal4/+ ;*

### Video 3.

*dnmnat*$^{OE}$: *w ; 20xUAS–IVS–CsChrimson::mVenus/5xUAS-dnmnat ; GMR60E02–Gal4/+ ;*
